# A Musashi-Related Protein is Essential for Gametogenesis in Arabidopsis

**DOI:** 10.1101/579714

**Authors:** Laura A. Moody, Ester Rabbinowitsch, Hugh G. Dickinson, Roxaana Clayton, David M. Emms, Jane A. Langdale

## Abstract

Musashi (Msi) proteins are an evolutionarily conserved group of RNA-binding proteins, required for targeted control of mRNA translation during many important developmental processes in animals. Most notably, Msi proteins play important roles during both spermatogenesis and oogenesis. Msi proteins also exist in plants but these are largely uncharacterized. Here we report the functional characterization of an Arabidopsis Msi ortholog *ABORTED GAMETOPHYTE 2* (*AOG2*), which encodes a protein containing two RNA recognition motifs and an ER-targeting signal. AOG2-GFP translational fusions were localized to the ER in transient assays, suggesting that AOG2 most likely binds to ER-targeted mRNAs. We show that disrupted *AOG2* function leads to a high rate of both ovule and seed abortion, and that homozygous loss of function mutants are embryo lethal. Furthermore, we demonstrate that *AOG2* is required to establish asymmetry during pollen mitosis I, and that loss of *AOG2* function leads to loss of pollen viability. Collectively the results reveal that AOG2 is required for the establishment of polarity and/or the progression of mitosis during gametophyte development in Arabidopsis, and thus Msi-related proteins have an evolutionarily conserved role in gametogenesis in both animals and plants.

**SIGNIFICANCE STATEMENT:** *ABORTED GAMETOPHYTE 2* (*AOG2*) encodes a Musashi-related RNA-binding protein that is required for gametogenesis and embryogenesis in Arabidopsis. *AOG2* is required for the establishment of polarity and/or the progression of mitosis during gametophyte development in Arabidopsis, and thus Musashi-related proteins have an evolutionarily conserved role in gametogenesis in both animals and plants.

## INTRODUCTION

Musashi (Msi) proteins belong to an evolutionarily conserved group of RNA-binding proteins, which are characterized by the presence of two RNA recognition motifs (RRMs) (Sutherland et al., 2013). Msi proteins are essential for the targeted control of mRNA translation, and many of the known Msi targets (e.g. Numb, a negative regulator of Notch signaling) play key roles in oncogenic pathways. Consequently, mis-regulation of Msi proteins is a major contributing factor in cancers and other diseases (Guo et al., 1995; Guo et al., 1996; Zhong et al., 1996; Lan et al., 2017; Zhang et al., 2014; Kharas et al., 2010; Ito et al., 2010; Griner et al., 2010; Kaeda et al., 2015; Li et al., 2015; Zong et al., 2016; Kudinov et al., 2016; Guo et al., 2017; Cox et al., 2013).

The Msi protein was originally identified as a neural RNA-binding protein that is essential for an asymmetric cell division in the Drosophila sensory organ. In WT, the sensory organ precursor (SOP) cell divides asymmetrically to give rise to a non-neuronal (IIa) and a neuronal (IIb) precursor cell. Socket and shaft cells of the external sensory organ of Drosophila are products of IIa cells whereas neurons and glia are products of IIb cells. In loss of *msi* mutants, the SOP undergoes an abnormal symmetric cell division to give rise to an overrepresentation of IIa cells, with no IIb cells. Consequently, sensory organs comprise only socket and shaft cells, and mutants exhibit a double bristle phenotype. This phenotype bears a resemblance to the two-sword fighting Japanese Samurai Miyamoto Musashi (Nakamura et al., 1994).

Msi proteins also play an important role during gametogenesis, and the role of Musashi-2 (Msi-2) in both spermatogenesis and oogenesis has been well described. For example, in mice, Msi-2 is highly expressed in meiotic spermatocytes and differentiating spermatids and is required to maintain a pool of spermatogonial stem cells (Siddall et al., 2006; Sutherland et al., 2014). Loss of Msi-2 function leads to premature differentiation of spermatids, and Msi-2 overexpression causes male sterility (Sutherland et al., 2015; Sutherland et al., 2018). During spermatogenesis, Msi-2 is required for the post-transcriptional regulation of both *Tbx1* and *Piwil1* mRNA (Sutherland et al., 2018). Msi-2 is also highly expressed during folliculogenesis and loss of Msi-2 function causes defective follicle development in the mouse ovary (Gunter and McLaughlin, 2011; Sutherland et al., 2015). Within Xenopus oocytes, Msi-2 binds to the 3’UTR of Mos mRNA during meiotic cell cycle progression and oocyte meiotic cell cycle progression is blocked when Msi-2 is prematurely truncated (Charlesworth et al., 2006).

Here we report the functional characterization of *ABORTED GAMETOPHYTE 2 (AOG2)*, a Msi-related protein in Arabidopsis. Loss of function mutant phenotypes reveal that AOG2 is required for both male and female gametophyte development, and for embryogenesis. Notably, asymmetric cell divisions are perturbed during pollen mitosis I, leading to loss of pollen viability in mutant plants. A role for Msi-related genes in both the establishment of asymmetry and the formation of gametes is thus conserved in animals and plants.

## RESULTS & DISCUSSION

### Eight Msi orthologs are present in the Arabidopsis genome

To identify *Msi2* orthologs in Arabidopsis, Orthofinder software (Emms & Kelly, 2015) was used to interrogate 11 genome sequences representing fungi, invertebrate and vertebrate animals, algae, plus non-flowering and flowering (both monocot and eudicot) plants. An orthogroup of 46 sequences was identified and aligned using Mergalign (Collingridge & Kelly, 2012) (Data S1-S3). Phylogenetic inference revealed an early duplication in the ophistokont lineage that generated separate *Msi* and *Daz-associated protein* (*Dazap*) clades, both of which are retained in the four animal genomes investigated (Figure 1). An early duplication is also inferred in the archeoplastida lineage. One of the resultant clades (‘plants I’) comprises single Arabidopsis (*At1g17640*) and moss (*Physcomitrella patens*) genes plus duplicated algal (*Chlamydomonas reinhardtii*) and monocot (*Oryza sativa, Sorghum bicolor*) genes. One of the *Msi* genes from *C. reinhardtii* (Cre12.g560300.t1.2) has previously been shown to play a role in thermal signalling but phylogenetic analyses in the same study did not identify the second (Cre07.g330300.t1.2) (Li et al., 2018). Although algal genes are not represented in the second clade (‘plants II’), multiple duplications are evident in each of the 5 plant genomes. Arabidopsis has seven genes in this clade-*At1g58470, At5g47620, At3g07810* plus recent duplicate pairs *At5g55550*/*At4g26650* and *At4g14300*/*At2g33410*.

**Figure 1.**
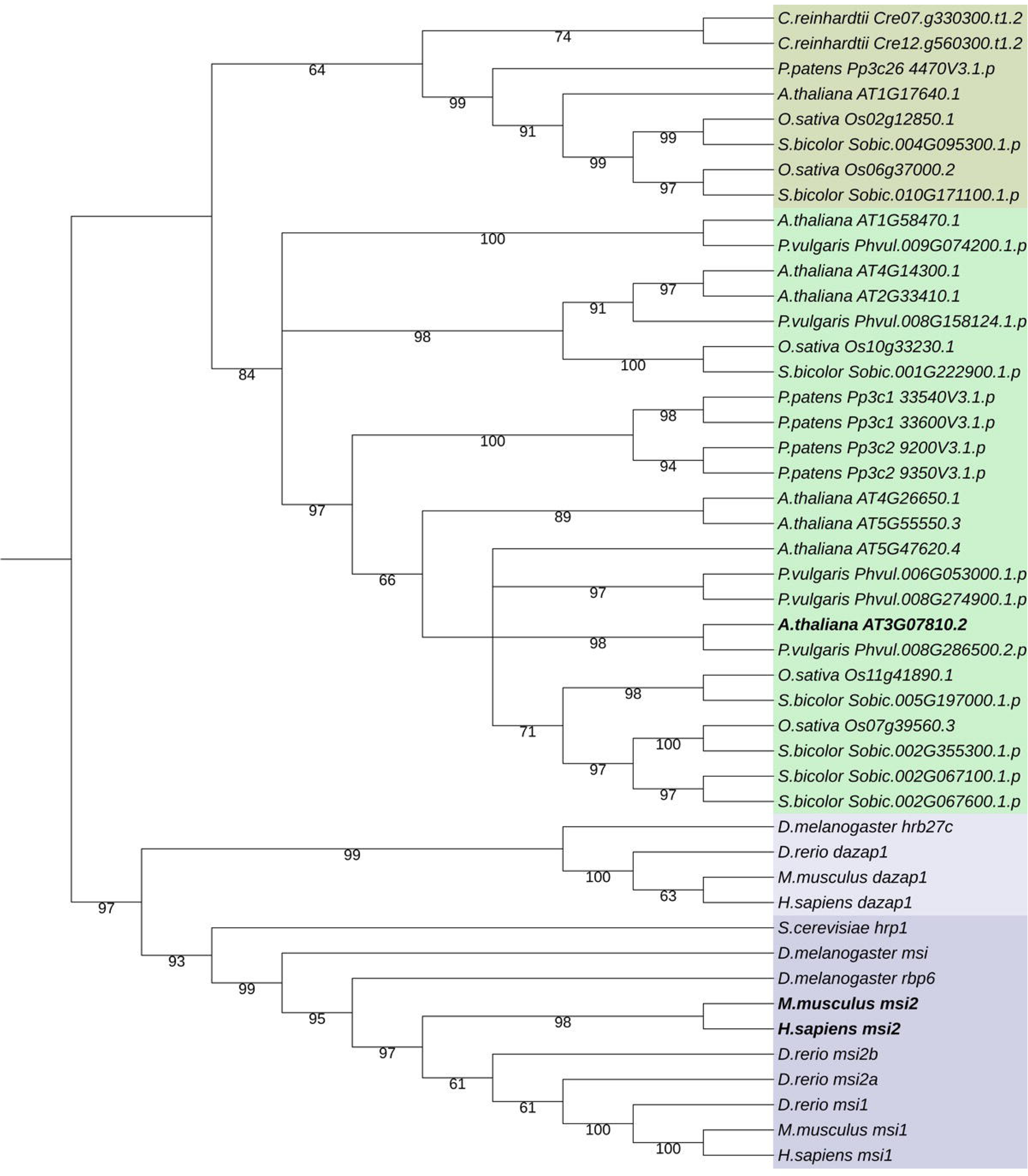
Phylogenetic analysis of Musashi homologs. Inferred phylogenetic tree from maximum likelihood analysis of 46 amino acid sequences retrieved from 11 genomes (see Data S1-S3 for accession numbers and alignment). Bootstrap values are given for each node. The tree was rooted on the branch between archeoplastida and ophistokonts and all branches with less than 50% bootstrap support were collapsed. Independent duplications in both groups resolved two clades in each (‘plants I’ - dark green; ‘plants II’ - light green; *Dazap* - light purple; *Msi* - dark purple).

Expression profiles of all seven Arabidopsis *Msi* orthologs in the ‘plants II’ clade were interrogated using GENEVESTIGATOR (Zimmermann et al., 2004). Transcripts of two of the genes (*At1g58470* and *At2g33410*) were enriched in the embryo, endosperm and seed coat. Four of the genes (*At5g47620, At3g07810, At5g55550, At4g26650*), which likely arose from within-eudicot or within-species duplications, were expressed most highly in pollen. One of the genes (*At4g14300)* was moderately expressed in both pollen and the embryo (Figure 2). The protein encoded by gene *At3g07810* is a Reciprocal Best Hit of metazoan MSI2 and vice versa, and as such is regarded as the closest MSI2 ortholog in Arabidopsis. To confirm that the protein encoded by the *At3g07810* gene accumulates in pollen, transgenic lines were generated in which 2.2kb of the promoter was used to drive expression of a translational fusion between the entire open reading frame (ORF) and the green fluorescent protein (GFP) reporter. The ORF-GFP fusion protein was diffusely localised to the cytoplasm of pollen grains (Figure S1). Based on these observations, we hypothesized that the Arabidopsis *Msi2* ortholog plays a role in pollen development.

**Figure 2.**
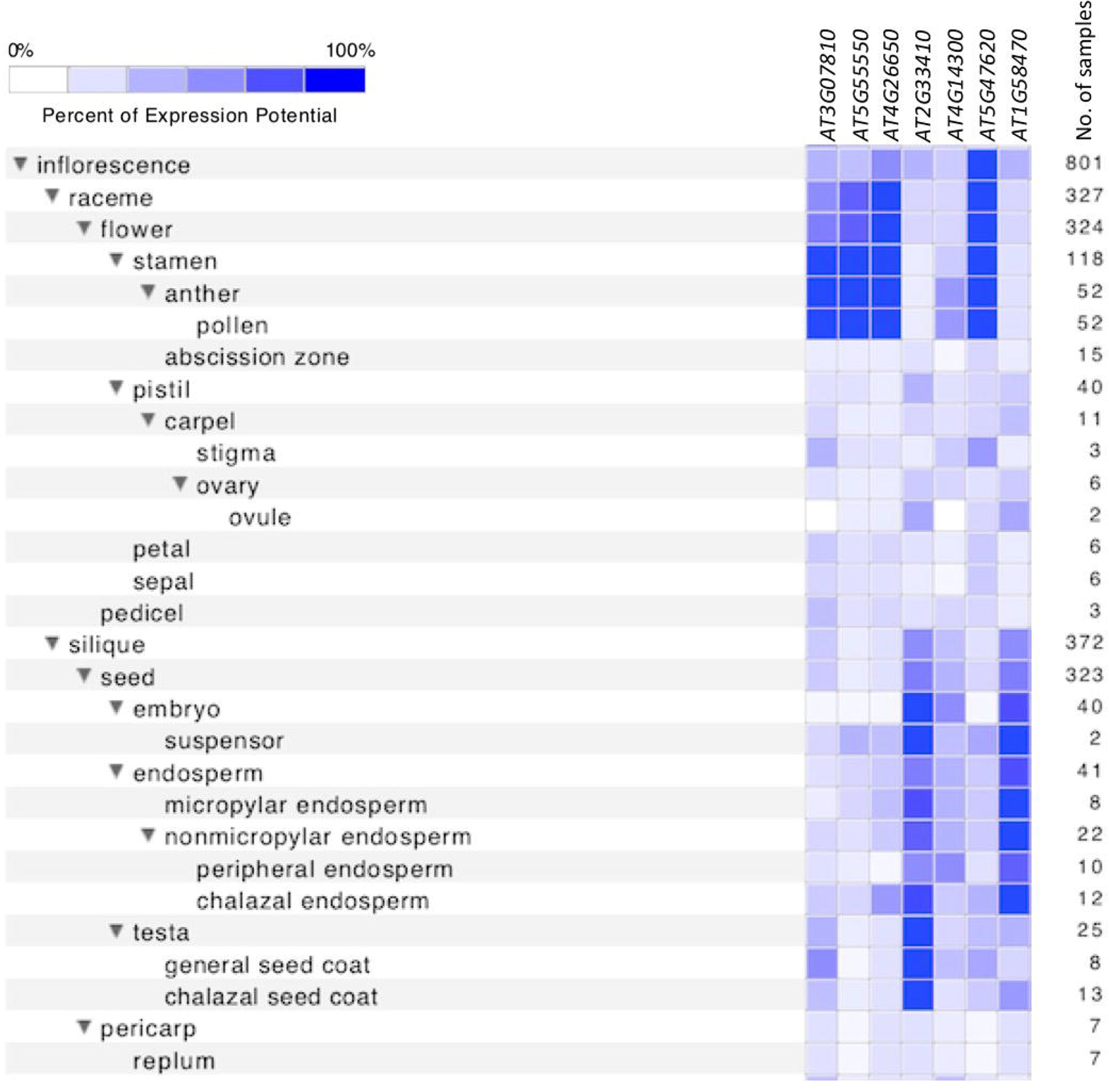
Expression of Arabidopsis Musashi-related genes. Summary of GENEVESTIGATOR ATH1 array data for the seven *Msi* orthologs in the ‘plants II’ clade. Dark blue boxes represent the highest signal intensity (100 %) whereas white boxes denote the absence of a signal.

### The Arabidopsis *Msi2* ortholog *AOG2 encodes a non-nuclear RNA-binding protein*

The *Msi2* ortholog *At3g07810*, which we named *ABORTED GAMETOPHYTE 2* (*AOG2*), encodes a protein of 495 amino acid residues with two RNA recognition motifs (RRM) at the N terminus, both of which are very similar to those in human *Msi2* (73.7% RRM1, 66.7% RRM2 (Figure 3a, Figure S2). An endoplasmic reticulum (ER) targeting signal was identified within RRM2 (using LocSigDB; Negi et al., 2015), and translational fusions with the reporter protein GFP were targeted to the ER in transient protoplast assays (Figure 3b). AOG2 therefore most likely binds to ER-localized RNAs in Arabidopsis and given the known function of animal orthologs most likely inhibits translation of the bound transcripts.

**Figure 3.**
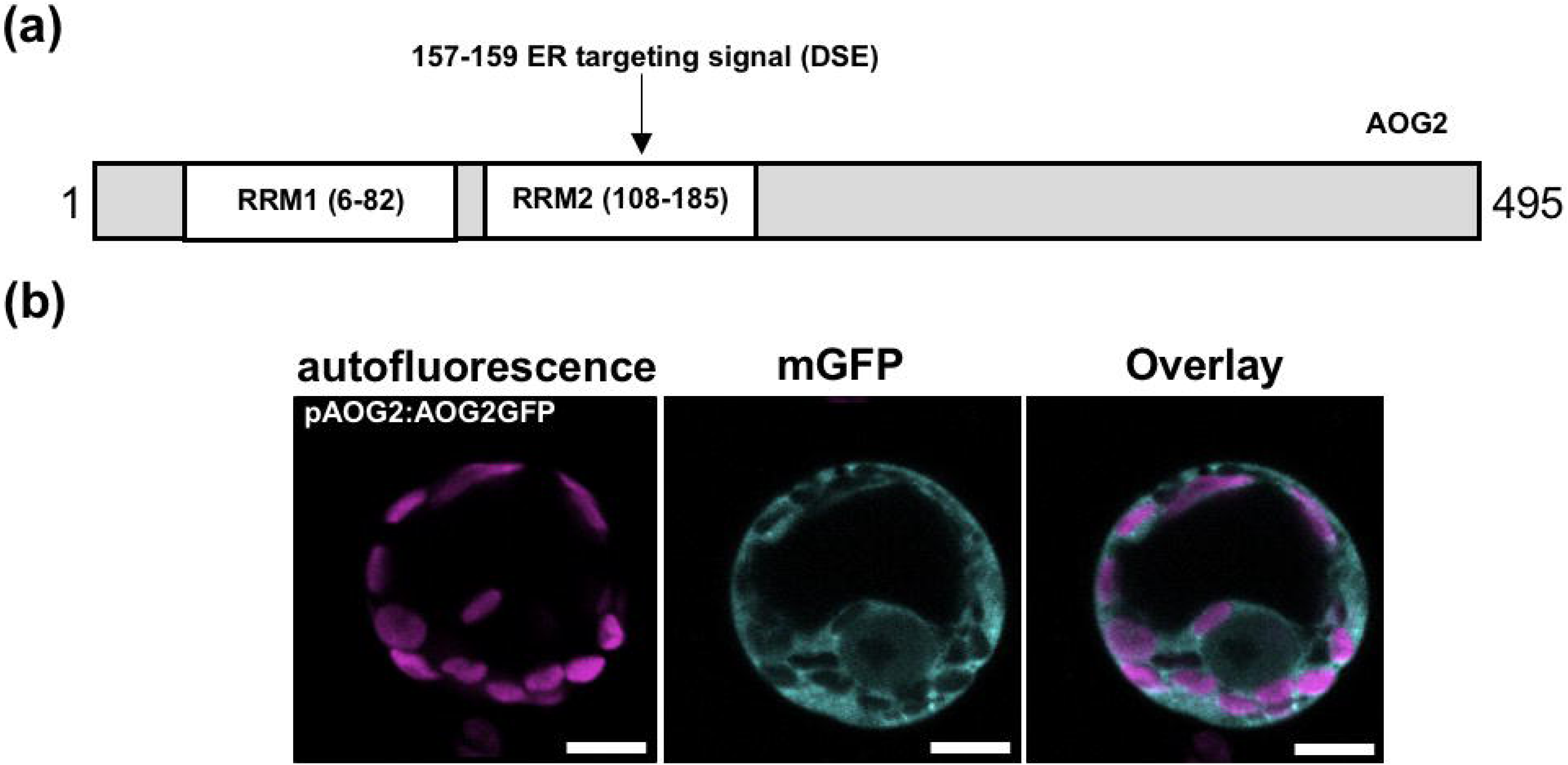
AOG2 localises to the endoplasmic reticulum. (a) The AOG2 protein (495 residues) contains two RNA recognition motifs (RRM1, 6-82; RRM2, 108-185) and an ER targeting signal (157-159). (b) AOG2-GFP localisation in protoplasts. Magenta represents autofluorescence, blue represents GFP and overlay represents merged autofluorescence and GFP images. Scale bar, 10 µm.

### Gametophyte development is perturbed in *aog2* mutants

To determine the function of *AOG2 in planta*, a T-DNA insertion line was obtained from the SALK collection (http://signal.salk.edu/cgi-bin/tdnaexpress) (Alonso et al., 2003). The SALK_007316 (*aog2-1*) line has a T-DNA insertion positioned between the open reading frames of two genes (*AOG2* and *At3g07800* which encodes the thymidine kinase AtTK1) (Figure 4a). The genomic arrangement is such that the T-DNA insertion could potentially influence the promoter activity of either or both of the flanking genes. However, it has previously been reported that null mutants of *AtTK1* are phenotypically indistinguishable from WT, and that gene function is associated with cell proliferation (Clausen et al., 2012, Xu et al., 2015). We thus concluded that any defects in pollen development in *aog2-1* mutants would be caused by reduced activity of *AOG2* rather than *AtTK1*, although the potential for synergistic effects could not be disregarded at this stage. Notably, *AOG2* expression levels were substantially reduced in *aog2-1* mutant pollen relative to WT (Figure 4b).

**Figure 4.**
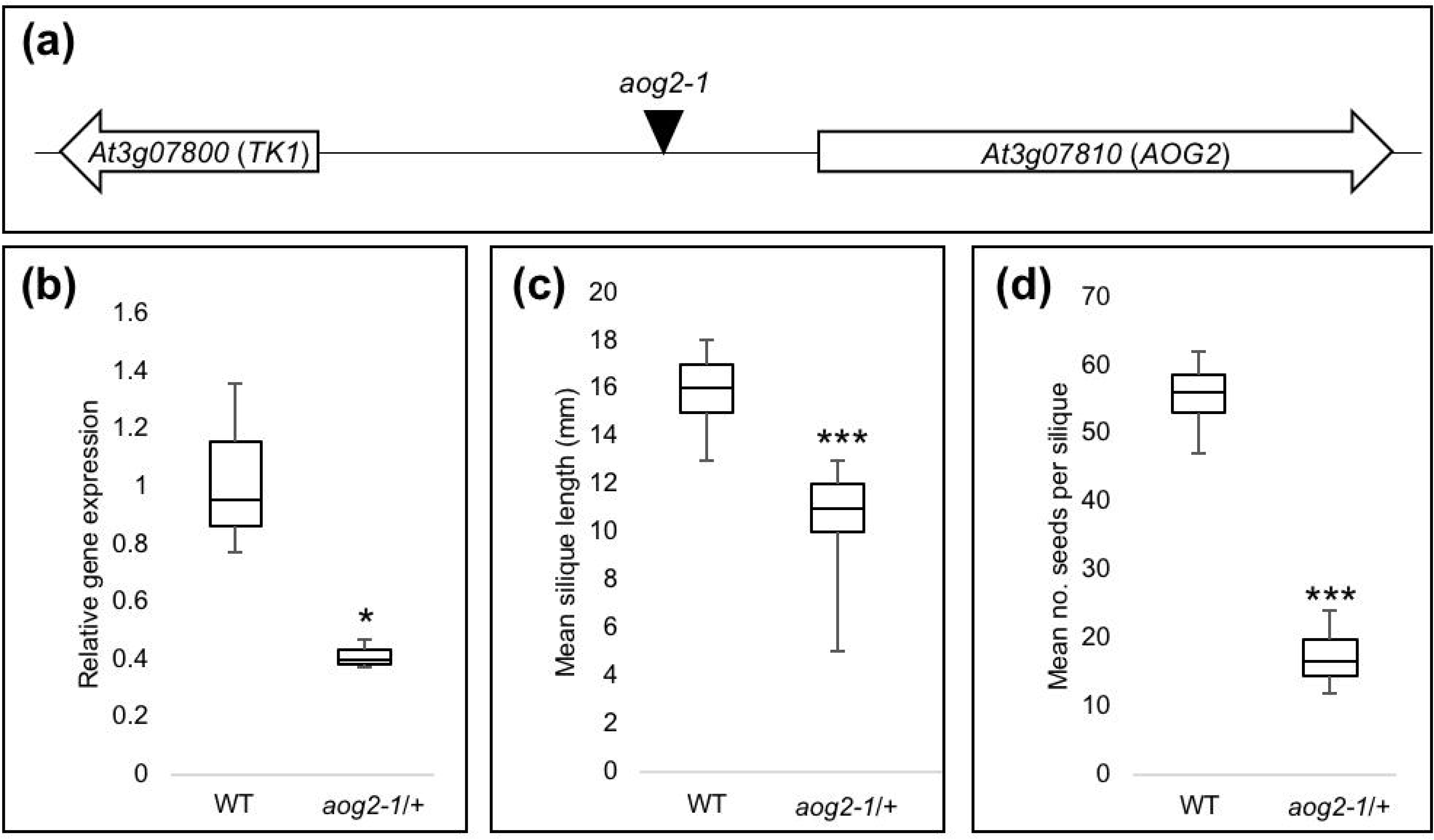
*aog2* mutants exhibit fertility defects. (a) T-DNA insertion (SALK_007316) positioned between the open reading frames of *AOG2* and *TK1* genes. (b) Quantitative RT-PCR data showing the relative expression of *AOG2* in pollen isolated from WT (Col-0) and *aog2-1*/*AOG2* plants. *t*-test, *, P<0.005. (c) Mean silique length in mature WT and *aog2-1*/*AOG2* plants. *t*-test, ***, P<0.001. (d) Mean number of seeds in WT and *aog2-1*/*AOG2* plants. *t*-test, ***, P<0.001.

Prior to phenotypic characterization, mutant lines were backcrossed to WT Col-0 plants. *aog2-1*/*AOG2* heterozygous plants were phenotypically indistinguishable from WT, except that mean silique lengths were significantly shorter (*aog2-1/AOG2* −11.40 ± 0.25 mm; WT −15.88 ± 0.46 mm) (Figure 4c) and the mean number of seeds per silique was significantly lower (*aog2-1/AOG2* −17.20 ± 1.28; WT - 55.3 ± 1.56) (Figure 4d). Given that reduced seed-set is observed in heterozygous plants, the *aog2-1* mutant allele either has a dominant effect in the diploid embryo or a recessive effect in the haploid gametophytes. Pollen-preferential *AOG2* expression in WT plants, reduced *AOG2* transcript levels in pollen from *aog2-1*/*AOG2* plants, and the observed seed-set phenotype are all consistent with a role for *AOG2* in Arabidopsis gametophyte development.

### Transmission through both male and female gametes is perturbed in *aog2* mutants

To test whether the reduced seed-set in *aog2-1*/*AOG2* heterozygotes resulted from compromised male (pollen) and/or female (megagametophyte) function, transmission efficiencies were quantified. The progeny of reciprocal test crosses between *aog2-1*/*AOG2* and WT plants were genotyped for the presence of the mutant *aog2-1* allele, and the transmission efficiency was deduced by calculating the percentage of *aog2-1*/*AOG2* individuals in the offspring. Only 32.6% of male gametes and 26.3% of female gametes carrying the *aog2-1* mutation were successfully transmitted to the next generation (Figure 5a). As such, *AOG2* is required for both male and female gametophyte development.

**Figure 5.**
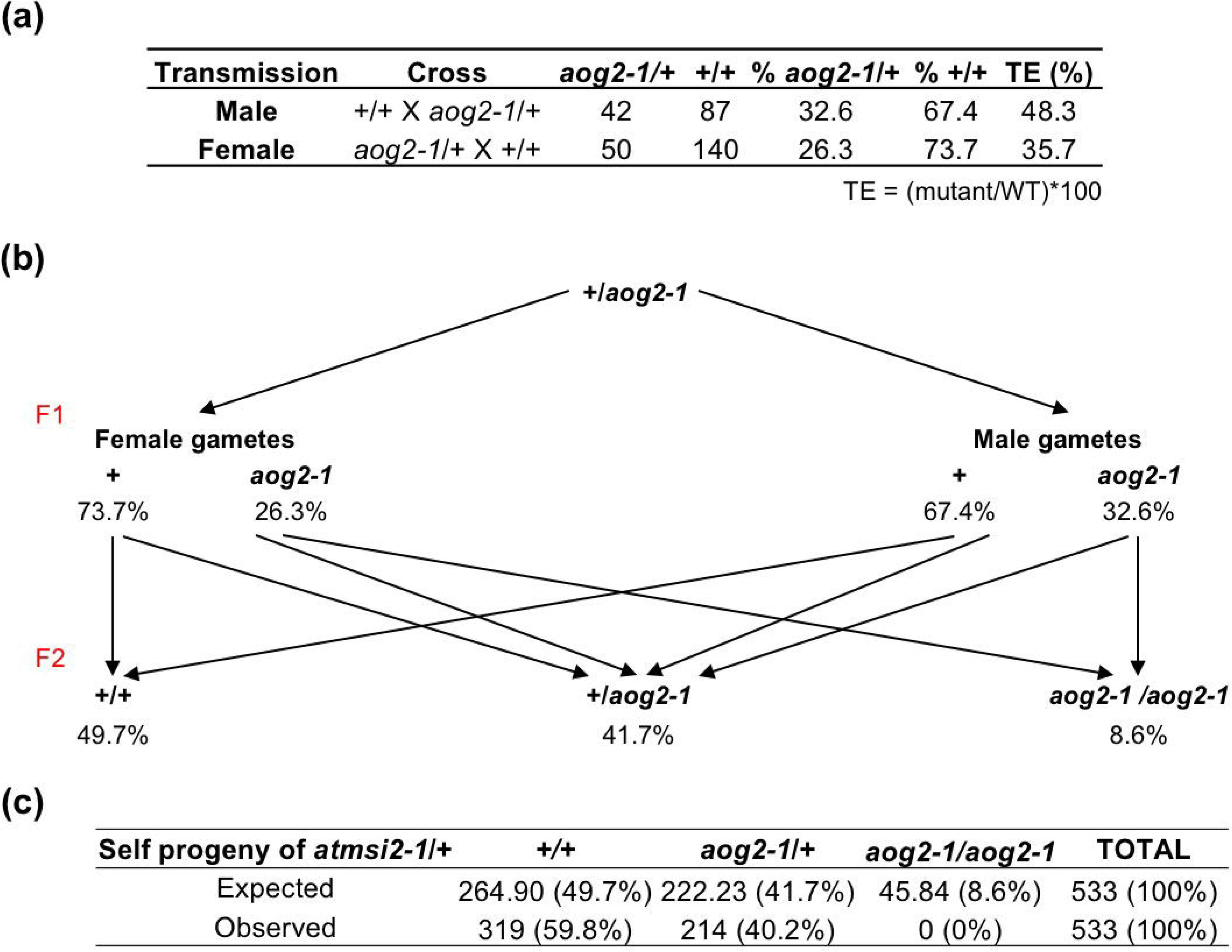
Genetic transmission of *aog2*. (a) The number of WT (+/+) and *aog2-1*/*AOG2* (aog2-1/+) individuals resulting from reciprocal test crosses between *aog2-1*/*AOG2* and WT (Col-0) plants. The transmission efficiency (TE) was calculated in each case. (b) Predicted frequencies of WT, *aog2-1*/+ and *aog2-1*/*aog2-1* progeny derived from self-progeny of *aog2-1*/+ plants. (c) Observed and expected numbers of WT, *aog2-1*/+ and *aog2-1*/*aog2-1* progeny derived from self-progeny of *aog2-1*/+ plants.

To characterize the homozygous mutant phenotype, F2 progeny of self-pollinated *aog2-1*/*AOG2* heterozygotes were analysed. Strikingly, no homozygous individuals were recovered in the offspring of *aog2-1*/*AOG2* plants. To determine whether defective transmission of the *aog2-1* allele was sufficient to account for this finding, the frequencies of WT, heterozygous and homozygous individuals amongst self-progeny of *aog2-1/AOG2* plants were predicted on the basis of the quantified transmission efficiencies (Figure 5b, 5c). In a genotyped population of over 500 F2 progeny, heterozygote frequency was as predicted (41.7% predicted versus 40.2% observed) and WT frequency was higher than expected (49.7% predicted versus 59.8% observed). Homozygous individuals were not identified despite a predicted frequency of 8.6% (i.e. 45 individuals in the population tested). This observation indicates that in addition to roles in gametophyte development, *AOG2* function may be required for development of the diploid embryo.

To confirm that loss of *AOG2* function was responsible for the gametophyte and embryo defects observed in mutant plants, *aog2-1*/*AOG2* plants were transformed with a construct in which 2.2kb of the *AOG2* promoter was used to drive expression of the *AOG2* coding sequence. In segregating T2 populations, homozygous *aog2-1/aog2-1* individuals that also contained the transgene were viable, confirming that the transgene complemented gametophyte defects. However, the silique length in these *aog2-1/aog2-1/AOG2* individuals was less than that seen in *aog2-1/AOG2* heterozygotes (Figure 6a-e). This observation suggests either that the silique phenotype is influenced by the proportion of mutant and WT alleles present, or that the regulatory sequences (promoter and/or UTRs) used in the transgene construct were not sufficient to replicate the spatial and temporal patterns of *AOG2* expression that are required to fully complement the *aog2-1* mutation.

**Figure 6.**
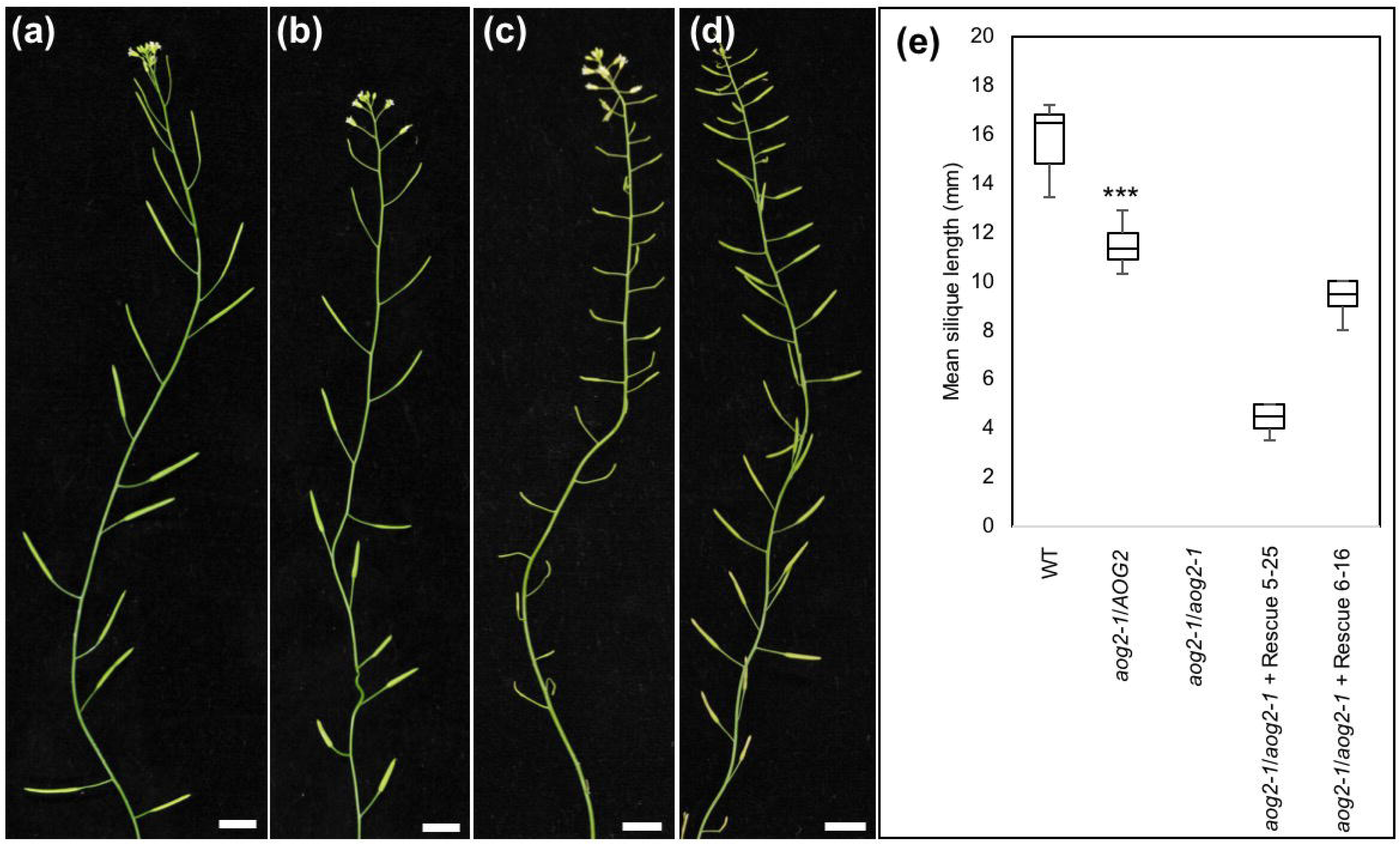
Complementation with full-length *AOG2* cDNA restores viability to *aog2-1*/*aog2-1* mutants. Representative images of (a) mature WT, (b) *aog2-1*/+, (c) *aog2-1*/*aog2-1* complementation line 5, (d) *aog2-1*/*aog2-1* complementation line 6. Scale bar, 10 mm. (e) Mean silique length in (from left to right) WT, *aog2-1*/*AOG2, aog2-1*/*aog2-1, aog2-1*/*aog2-1* complementation line 5 and *aog2-1*/*aog2-1* complementation line 6. *t*-test, ***, P<0.001.

### *AOG2* is required for both female gametophyte and embryo development

To investigate how loss of *AOG2* function specifically affects female gametophyte development, siliques from *aog2-1*/*AOG2* plants were dissected. Notably, a significant proportion (approximately 50%) of ovules aborted prior to fertilization as evidenced by empty spaces in the ovary (Figure 7a-d). In addition, a high rate of seed abortion was observed in siliques derived from *aog2-1*/*AOG2* (10.81 ± 2.71) plants compared to WT (1.03 ± 0.38) (Figure 7e). Mature siliques of *aog2-1*/*AOG2* heterozygous plants thus contained a mixture of fully developed and embryo-arrested seeds. Developmental arrest was observed as early as the 2-cell stage (Figure 7f) but was also observed at the late globular (Figure 7g) and late heart stages (Figure 7h). Collectively these observations confirm a role for *AOG2* both in female gametophyte and embryo development.

**Figure 7.**
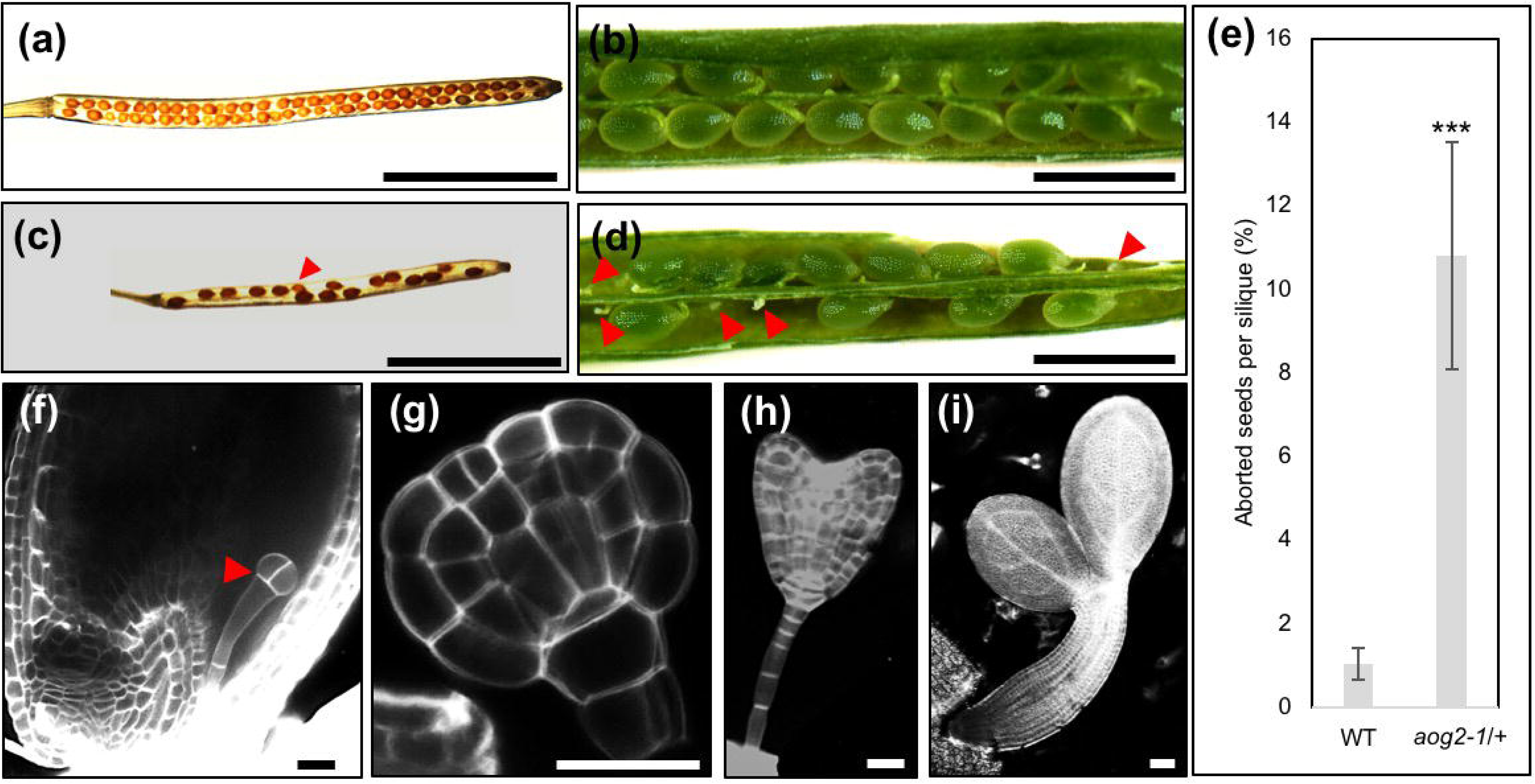
*AOG2* is essential for female gametophyte development. (a, c) Cleared siliques from WT and *aog2-1*/+ respectively. Scale bar, 5 mm. Aborted seeds denoted by an arrow in (c). (b, d) Dissected siliques from WT and *aog2-1*/+ respectively. Scale bar, 1 mm. Aborted ovules denoted by red arrows in (d). (e) Percentage of aborted seeds per silique in WT and *aog2-1*/+. *t*-test, ***, P<0.001. (f-i) Developmental arrest observed at the two-cell stage (f), globular stage (g) and late heart (h) of embryos in *aog2-1*/+. Normal embryo development was otherwise observed (i).

### *AOG2* is required for pollen mitosis I

To determine the role of *AOG2* in male gametophyte development, pollen viability was first observed using Alexander staining, which stains the cytoplasm of viable pollen grains red and that of non-viable or aborted grains green (Figure 8a-c). Quantification revealed that the mean number of viable mature pollen grains per anther was significantly reduced in *aog2-1/AOG2* plants (87.83 ± 10.80) compared to WT (213.44 ± 11.72) (Figure 8d), consistent with the observed transmission efficiency of 48% (Figure 5a).

**Figure 8.**
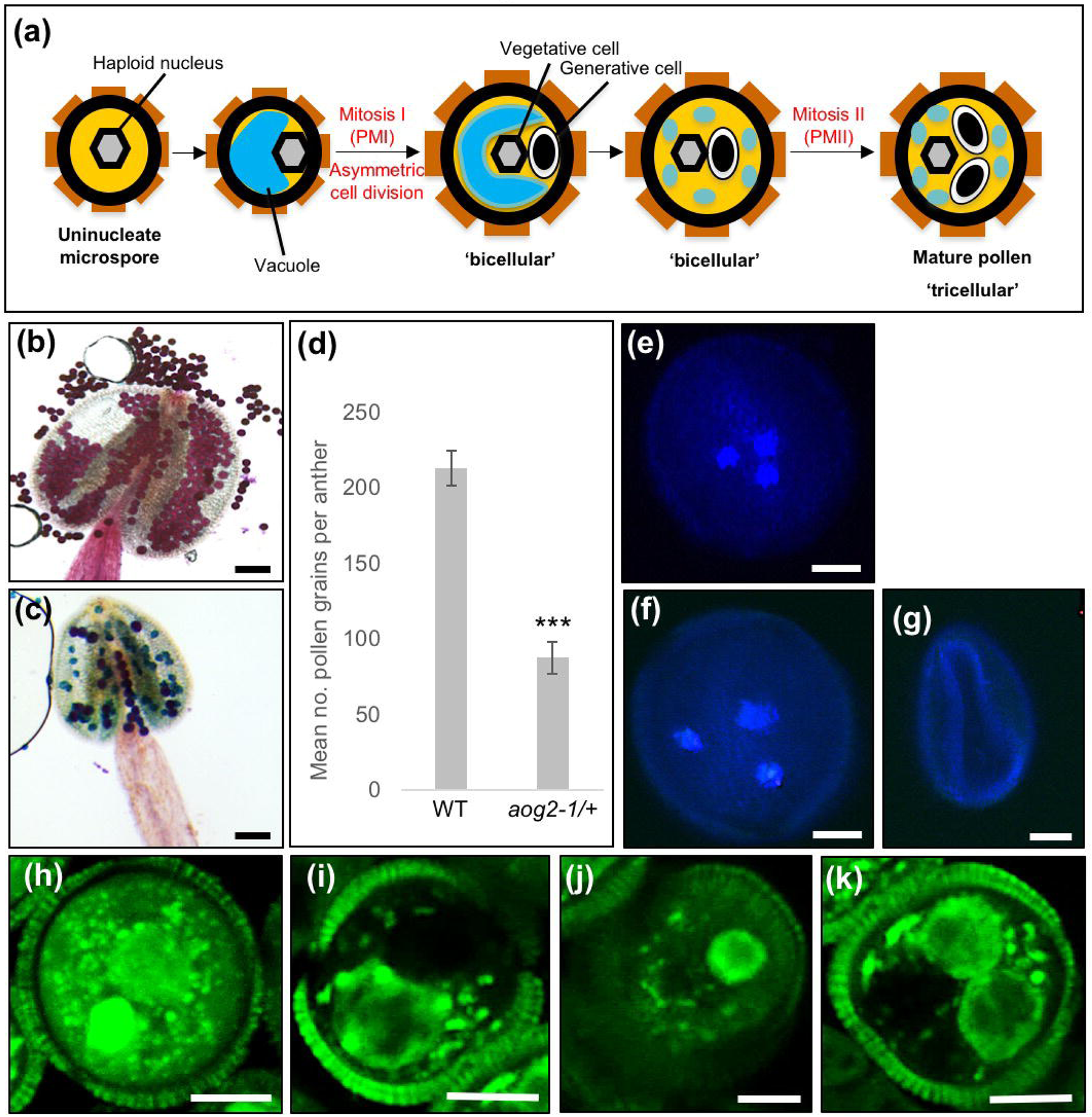
*AOG2* is essential for male gametophyte development. (a) Diagram depicting the stages of pollen development. PMI and PMII denote Pollen Mitosis I and Pollen Mitosis II respectively. (b, c) Alexander-stained anthers from WT (b) and *aog2-1*/+ (c) containing viable (red) and non-viable or aborted pollen grains (green). Scale bar, 100 µm. (d) The mean number of pollen grains per anther in WT and *aog2-1*/+. *t*-test, ***, P<0.001. (e-g) Normal tricellular pollen in WT (e) and an equal mixture of tricellular pollen (f) and aborted unicellular microspores (g) in *aog2-1*/+. (h-k) DAPI-stained cryostat-sectioned anthers from WT (h) and aog2-1/+ (i-k). Abnormal asymmetric cell division shown in (k). Scale bar, 5 µm.

To understand how the *aog2* mutation affects pollen development, DAPI-stained mature pollen grains were examined by confocal microscopy. In WT plants, the development of pollen grains begins with the formation of diploid mother cells (i.e. meiocytes), which divide meiotically to form four haploid unicellular microspores. Each microspore subsequently divides asymmetrically during pollen mitosis I (PMI) to produce two unequal daughter cells; a vegetative cell and a generative cell. The vegetative cell adopts a terminal cell fate whereas the generative cell divides once more during pollen mitosis II to produce two sperm cells, enclosed within the cytoplasm of the vegetative cell. Mature pollen that is released from the anthers contains three cells (tricellular pollen) (Figure 8a). In WT DAPI-stained pollen, tricellular pollen was invariably detected in which one vegetative nucleus and two sperm nuclei were easily distinguishable (Figure 8e). However, in *aog2-1*/*AOG2* DAPI-stained pollen, a roughly equal mixture of both tricellular pollen (Figure 8f) and aborted unicellular microspores (Figure 8g) were detected. Loss of *AOG2* function thus prevented pollen mitosis I.

To confirm that *aog2-1* pollen is unable to undergo normal pollen mitosis, cryostat-sectioned anthers were DAPI stained. In samples from *aog2-1*/*AOG2* plants, normal PMI was observed in approximately half of the microspores observed but in the other half a symmetric cell division gave rise to two nuclei of unknown fate (Figure 8h-k). Microspores that exhibited defects in PMI did not proceed beyond the vacuolar microspore stage. Collectively these results demonstrate that *AOG2* is required for the asymmetric cell division at PMI that produces the vegetative and generative cells, and that loss of function leads to loss of pollen viability.

## Conclusion

The *aog2-1* mutant phenotypes, namely disrupted asymmetric cell division during PMI, reduced transmission efficiencies of both male and female gametes, and perturbed embryo development are similar to those reported for mutations in *GEMINI POLLEN 1* (*GEM1* - also known as *MOR1*) (Park et al., 1998; Park and Twell, 2001; Whittington et al., 2001; Twell et al., 2002), *GEM2* (Park et al., 2004) and *GEM3* (also known as *AUGMIN SUBUNIT 6*) (Oh et al., 2016). *GEM1* encodes a microtubule-associated protein and *GEM3* is required for microtubule organisation and mitotic progression. Additional proteins that are essential for gametophyte development include ABORTED GAMETOPHYTE 1 (Cui et al., 2015), the DC1-domain protein VACUOLELESS GAMETOPHYTES (D’Ippólito et al., 2017), and the leucine-rich repeat protein PIRL6 (Forsthoefel et al., 2018). Whether AOG2 acts upstream or downstream of these other factors remains to be determined, but if any of them are translated in the ER, they are putative AOG2 targets.

## EXPERIMENTAL PROCEDURES

### Phylogeny construction

Genome sequences of yeast (*Saccharomyces cerevisiae*), fruitfly (*Drosophila melanogaster*), zebrafish (*Danio rerio*), mouse (*Mus musculus*) and human (*Homo sapiens*) were extracted from ENSEMBL (www.ensembl.org/index.html), and those of the alga *Chlamydomonas reinhardtii*, the moss *Physcomitrella patens*, the eudicots thalecress (*Arabidopsis thaliana*) and bean (*Phaseolus vulgaris*), plus the monocots rice (*Oryza sativa*) and sorghum (*Sorghum bicolor*) were extracted from Phytozome (www.phytozome.jgi.doe.gov/pz/portal.html). 99 sequences comprising the orthogroup that includes the mouse *msi2* gene were retrieved using Orthofinder (Emms and Kelly, 2015) (Data S1) and aligned using Mergalign (Collingridge and Kelly, 2012) (Data S2). The alignment was manually trimmed using Mega7 (Kumar et al. 2016) to remove highly variable regions. Where genes were represented by multiple transcript variants, all but the longest variant was removed from the dataset. The zebrafish CR931806.1 and *msi* homolog-like sequences were also removed because both were truncated. The trimmed alignment comprised 46 sequences (Data S3). The best-fitting model parameters (LG+I+G4) were estimated and consensus phylogenetic trees were run using Maximum Likelihood from 1000 bootstrap replicates, using IQTREE (Nguyen et al. 2015). The data were imported into ITOL (Letunic and Bork, 2016) to generate the pictorial representation. The tree was rooted on the branch between ophistokonts and archeoplastida, and all branches with less than 50% bootstrap support were collapsed.

### Plant strains and growth conditions

Seeds were grown in Levington M3 compost in the greenhouse at 22°C with a 16 h light (300µmol photons m^-2^s^-1^): 8 h dark cycle. The segregating, single-insert *aog2-1* (SALK_007316) line was obtained from the Nottingham Arabidopsis Stock Centre. Segregating *aog2-1* lines were genotyped using AOG2_LP and AOG2_RP primers to detect the WT allele), and LBa1 and AOG2_RP primers to detect the mutant allele. Primers were designed using the T-DNA Primer Design tool (SIGnAL). The primers AOG2_LP and AtMSI2_intron_2R were used to detect the presence of the WT allele in rescue lines.

### Reciprocal crosses

To determine transmission of the mutation through the female, *aog2-1*/*AOG2* mutant plants were cross-pollinated with WT pollen. To determine transmission of the mutation through the male, WT plants were cross-pollinated with *aog2-1*/*AOG2* pollen. Progeny of crosses were genotyped by PCR to check for genetic segregation.

### RNA isolation and quantitative PCR

Total RNA was prepared from pollen grains using the RNeasy plant kit (Qiagen). 1 µg RNA was DNase treated (TURBO DNase, Ambion) and then cDNA was synthesised using Superscript III reverse transcriptase (Invitrogen).

The expression of *AOG2* was analysed by quantitative real-time Reverse Transcription PCR (qRT-PCR). Primers were designed to amplify a 100-150 bp fragment of each gene (Primer 3). Amplification was detected using the SYBR Green master mix (Life Technologies) according to the manufacturer’s specifications on a StepOne™ Real-Time PCR System (Applied Biosystems). Cycling conditions were 95°C for 5 min, and 40 cycles of 95°C for 15 s and 60°C for 1 min. Three technical replicates were performed for each of three independent biological samples, alongside water controls. Ct values were calculated from raw amplification data using the Real-time PCR Miner software (http://www.ewindup.info/miner/). The mean Ct value between the three technical replicates was then calculated. Fold changes in gene expression were calculated relative to controls using the 2^^ (-ΔΔCt)^ method. Two housekeeping genes were used as constitutive controls; *EF1B* (At5g19510) and *UBP6* (At1g51710). To test for genomic DNA contamination, no RT controls were run to check that no amplification had occurred.

### Cloning and construct generation

To generate the *AOG2* rescue construct, the *AOG2* promoter sequence was amplified from WT Col-O genomic DNA using pAOG2FSalI and pAOG2RBamHI and inserted into SalI/BamHI cut pBJ36 (Eshed et al., 2001) to create *AOG2pro*-*ocs3’*. The full-length *AOG2* coding region was amplified from cDNA prepared from flower mRNA using AOG2StartBamHI and AOG2RinclStopXbaI, and inserted into BamHI/XbaI cut *AOG2pro*-*ocs3’* to create *AOG2pro*:*AOG2*-*ocs3’*. The *AOG2pro*:*AOG2*-*ocs3’* cassette was transferred as a NotI fragment into the binary vector pMLBART (Eshed et al., 2001) and transformed into *aog2-1*/*AOG2* plants by floral dipping (Clough and Bent, 1998). Transformed plants were selected on the basis of Basta resistance and presence of the T-DNA insertion.

To generate the *AOG2pro*:AOG2-GFP reporter construct, the *AOG2* promoter sequence was amplified from WT Col-O genomic DNA using pAOG2FSalI and pAOG2RBamHI and inserted into SalI/BamHI cut pGreen (Hellens et al., 2000) to create *AOG2pro*-GFPter. The full-length *AOG2* coding region (without the stop codon) was amplified from cDNA prepared from flower mRNA using AOG2StartBamHI and AOG2-StopNotI and inserted into BamHI/NotI cut *AOG2pro*-GFPter to create *AOG2pro*::*AOG2*-GFPter.

### Transient assays protoplasts

10µg *AOG2pro*:AOG2-GFP reporter construct was transiently transformed into protoplasts isolated from *Physcomitrella patens* as described in Moody et al. (2012).

### Silique measurements and seed counts

Length measurements and seed counts were carried out for ten siliques derived from the primary inflorescence of each of three plants and repeated three times. Mature siliques were harvested and cleared in clearing solution (0.2 M NaOH, 1 % SDS).

### Isolation of mature pollen grains

Mature pollen grains were isolated from stage 13 flowers (Sanders et al., 1999). Flowers from approximately ten plants were collected and placed into 500 ml flasks containing 100 ml ice-cold 0.3 M mannitol, and then vigorously shaken by hand for approximately 3 minutes. Pollen suspensions were then filtered through a 100 µm nylon mesh into 50 ml falcon tubes. Pollen grains were harvested by centrifugation at 5000xg for 15 minutes. This step was repeated until all the pollen suspension had been used. After the final centrifugation, 2 ml of 0.3 M mannitol was left in the bottom of the tube and used to resuspend the pollen pellet and transfer it to a 2 ml eppendorf tube. The pollen grains were collected by centrifugation at maximum speed for 1 minute and then frozen in liquid nitrogen.

### DAPI staining

Pollen grains were harvested by centrifugation and resuspended in phosphate buffered saline (PBS) containing 2 µg/ml DAPI. Pollen grains were incubated overnight at room temperature in the dark. DAPI stained pollen grains were then imaged in a drop of water using a confocal microscope.

### Calcofluor staining

Embryos and seed coats were separated using fine tweezers. Embryos were then submerged in a 10 µg/ml calcofluor solution for 5 min, rinsed with water and then visualised using confocal microscopy.

### Alexander staining and determination of pollen viability

Anthers were removed from mature flowers (stage 13, Sanders et al., 1999) and placed onto a microscope slide. A drop of Alexander staining solution was added using a Pasteur pipette. Alexander staining solution was prepared as previously described (Peterson et al., 2010). Stained pollen grains were observed using a Leica M165C microscope equipped with a QImaging Micropublishing 5.0 RTV camera. Two anthers were selected for each of three flowers per plant (three plants in total).

### Microscopy

Fluorescence microscopy was carried out with a Zeiss LSM510 META confocal microscope. A 40x water immersive lens (C-Apochromat 40x/1.20 W) was used for all imaging. GFP was excited with the 458 nm laser line and detected with a 475-525 nm bandpass filter. Calcofluor was excited at 405 nm and detected with a LP420 nm filter. Other images were captured using either a Leica DMRB or a Leica M165C microscope, equipped with a QImaging Micropublishing 5.0 RTV camera.

### Cryosectioning

Detached inflorescences of WT Col-0 and heterozygous *aog2-1*/*AOG2* lines were fixed at room temperature for 3 h in 4 % paraformaldehyde in 0.05 M PHEM buffer (Brown and Lemmon, 1995), pH 7.0, containing 5 % DMSO and 0.01 % TRITON-X100. Following fixation, the material was washed in buffer for 15 min and equilibrated for 2 h each in successive solutions of 7 % and 12 % sucrose in the same buffer/DMSO/TRITON combination. Inflorescences were then transferred to a drop of OCT Tissue Plus embedding medium (Scigen, CA, USA) on a slide, all bubbles removed using a dissecting needle, and then snap-frozen in fresh OCT Tissue Plus in a thin-walled plastic embedding mould floating in solid CO_2_ cooled pentane. The frozen blocks were sectioned at 10 µm on a Reichert-Jung 2800 Frigocut cryostat and sections fixed onto Superfrost slides (Menzel, Saabruken, Germany) with a thin film of agar/gelatine (Brown and Lemmon, 1995). Sections were stained for 2 h in PBS containing 2 µg/ml DAPI and mounted in a 1:1 mix of glycerol and water containing 1 % DABCO antifade (Sigma-Aldrich, USA) for confocal imaging using a Zeiss (Cambridge, UK) CSLM 410 (Brown and Lemmon, 1995).

## ACCESSION NUMBERS

Accession numbers for genes described in this article are At3g07810 (AOG2) and At3g07800 (TK1). These can easily be found at either TAIR (https://www.arabidopsis.org) or Phytozome (https://phytozome.jgi.doe.gov/pz/portal.html).

## Supporting information

Primer Table

## ACKNOWLEDGEMENTS

We are grateful to Juliet C. Coates for providing the pGreen vector. The work was funded by a BBSRC (BB/M020517/1) grant to JAL.

## CONFLICT OF INTEREST

We declare no conflict of interest.

## SUPPORTING INFORMATION

**Data S1.** Sequences comprising the orthogroup that includes Msi2

**Data S2.** Alignment of Msi2-related proteins (Mergalign)

**Data S3.** Trimmed alignment of Msi2-related proteins (46 sequences)

## AUTHOR CONTRIBUTIONS

L.A.M. and J.A.L. conceived and designed the study; L.A.M. and J.A.L. wrote the manuscript with input from H.G.D.; E.R. performed reciprocal crosses; H.G.D. carried out the cryosectioning; J.A.L. performed phylogenetic analyses, assisted by D.M.E; L.A.M performed all other experimental work, with technical assistance (e.g. genotyping) from E.R. and R.C.

---

**Figure S1.**
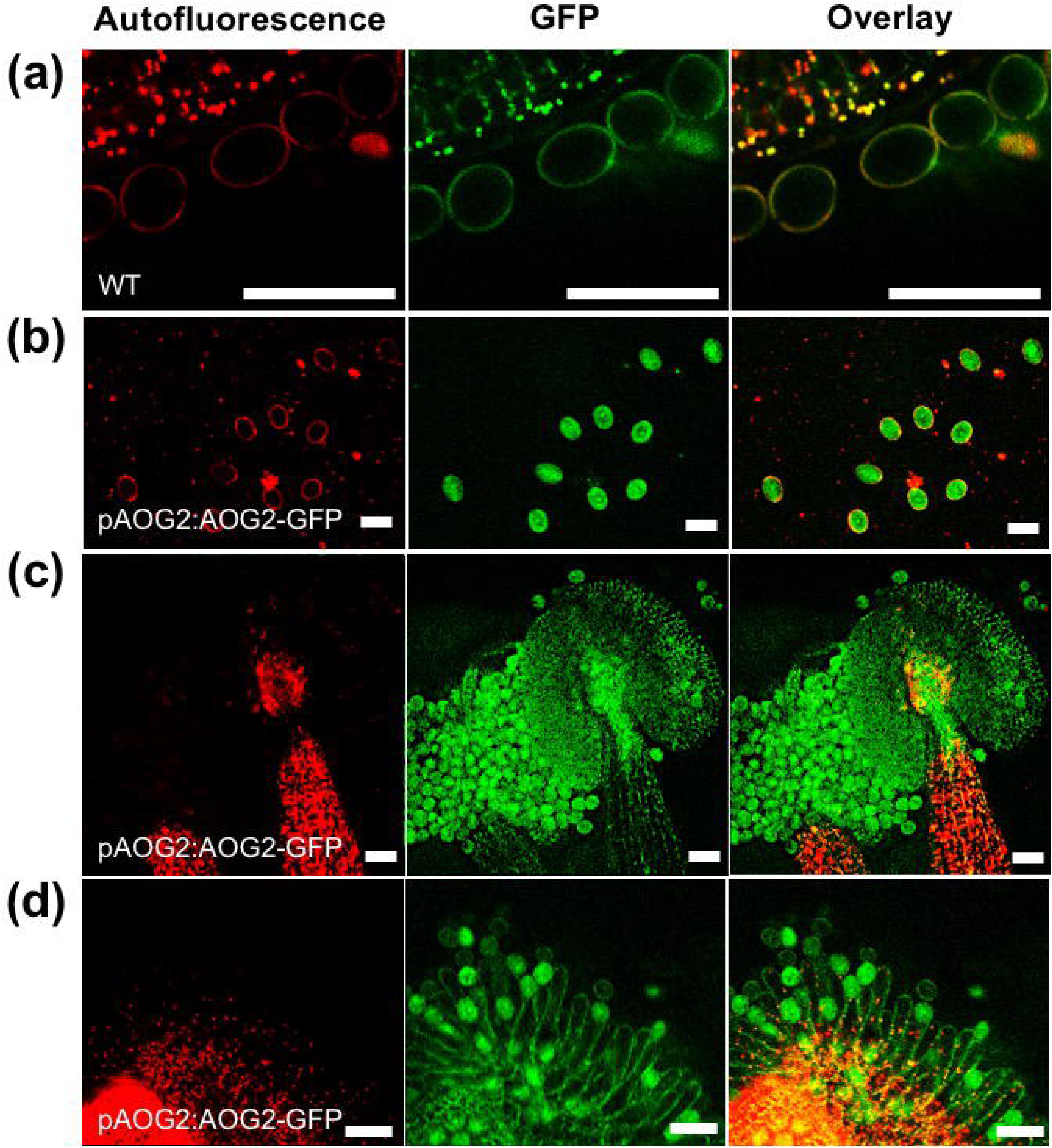
AOG2 is expressed in mature pollen grains. (a) Autofluorescence in WT pollen grains. (b-d) pAOG2:AOG2-GFP expression in pollen grains (b), in pollen grains within the anther (c) and in pollen grains on stigma (d). Red represents autofluorescence, green represents GFP and overlay represents merged autofluorescence and GFP images. Scale bar, 50 µm.

**Figure S2.**
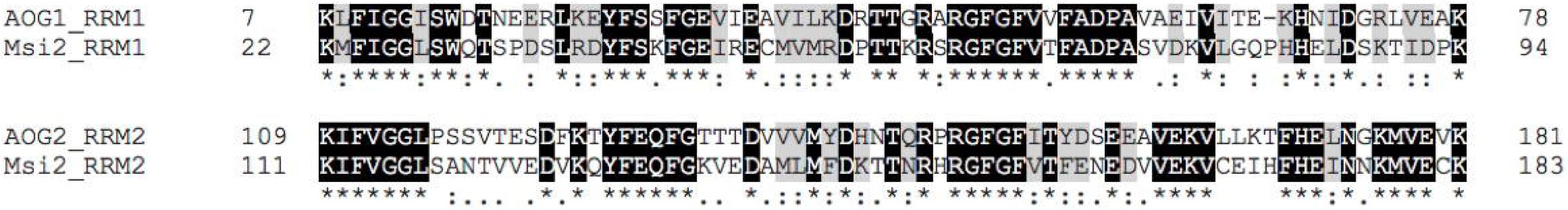
The AOG2 RRMs are similar to those from human Msi2. An alignment of both RRM1 and RRM2 from AOG2 and human Msi2. Identical amino acids are denoted by an asterisk (*), conserved amino acids denoted by a colon (:) and semi-conserved amino acids by a period (.). Numbers to the left and right of the alignment correspond to the relative position of amino acids from the start residue, methionine.

