## Supplementary material for "A Musashi-Related Protein is Essential for Gametogenesis in Arabidopsis": Data S1

>Athaliana_167_TAIR10.protein_primaryTranscriptOnly.fa_AT2G33410.1

MESDQGKLFIGGISWDTDENLLREYFSNFGEVLQVTVMREKATGRPRGFGFVAFSDPAVIDRVLQDKHHIDNRDVDVKRA

MSREEQSPAGRSGTFNASRNFDSGANVRTKKIFVGGLPPALTSDEFRAYFETYGPVSDAVIMIDQTTQRPRGFGFVSFDS

EDSVDLVLHKTFHDLNGKQVEVKRALPKDANPGIASGGGRGSGGAGGFPGYGGSGGSGYEGRVDSNRYMQPQNTGSGYPP

YGGSGYGTGYGYGSNGVGYGGFGGYGNPAGAPYGNPSVPGAGFGSGPRSSWGAQAPSGYGNVGYGNAAPWGGSGGPGSAV

MGQAGASAGYGSQGYGYGGNDSSYGTPSAYGAVGGRSGNMPNNHGGGGYADALDGSGGYGNHQGNNGQAGYGGGYGSGRQ

AQQQ*

>Athaliana_167_TAIR10.protein_primaryTranscriptOnly.fa_AT4G14300.1

MDSDQGKLFVGGISWETDEDKLREHFTNYGEVSQAIVMRDKLTGRPRGFGFVIFSDPSVLDRVLQEKHSIDTREVDVKRA

MSREEQQVSGRTGNLNTSRSSGGDAYNKTKKIFVGGLPPTLTDEEFRQYFEVYGPVTDVAIMYDQATNRPRGFGFVSFDS

EDAVDSVLHKTFHDLSGKQVEVKRALPKDANPGGGGRSMGGGGSGGYQGYGGNESSYDGRMDSNRFLQHQSVGNGLPSYG

SSGYGAGYGNGSNGAGYGAYGGYTGSAGGYGAGATAGYGATNIPGAGYGSSTGVAPRNSWDTPASSGYGNPGYGSGAAHS

GYGVPGAAPPTQSPSGYSNQGYGYGGYSGSDSGYGNQAAYGVVGGRPSGGGSNNPGSGGYMGGGYGDGSWRSDPSQGYGG

GYNDGQGRQGQ*

>Athaliana_167_TAIR10.protein_primaryTranscriptOnly.fa_AT4G26650.1

MNPEEQKMESASDLGKLFIGGISWDTDEERLQEYFGKYGDLVEAVIMRDRTTGRARGFGFIVFADPSVAERVIMDKHIID

GRTVEAKKAVPRDDQQVLKRHASPMHLISPSHGGNGGGARTKKIFVGGLPSSITEAEFKNYFDQFGTIADVVVMYDHNTQ

RPRGFGFITFDSEESVDMVLHKTFHELNGKMVEVKRAVPKELSSTTPNRSPLIGYGNNYGVVPNRSSANSYFNSFPPGYN

NNNLGSAGRFSPIGSGRNAFSSFGLGLNQELNLNSNFDGNTLGYSRIPGNQYFNSASPNRYNSPIGYNRGDSAYNPSNRD

LWGNRSDSSGPGWNLGVSVGNNRGNWGLSSVVSDNNGYGRSYGAGSGLSGLSFAGNTNGFDGSIGELYRGSSVYSDSTWQ

QSMPHHQSSNELDGLSRSYGFGIDNVGSDPSANASEGYSGNYNVGNRQTHRGIEA*

>Athaliana_167_TAIR10.protein_primaryTranscriptOnly.fa_AT1G17640.1

MDYNSDRGYDDSYHHHVHDEQLPQSDFEVETFDDRRNGGAAVDTGGIQMKHSVDHRHSSSSMSSPGKLFVGGVSWETTAE

TFANYFGKFGEVVDSVIMTDRITGNPRGFGFVTFADSAVAEKVLEEDHVIDDRKVDLKRTLPRGDKDTDIKAVSKTRKIF

VGGLPPLLEEDELKNYFCVYGDIIEHQIMYDHHTGRSRGFGFVTFQTEDSVDRLFSDGKVHELGDKQVEIKRAEPKRTGR

DNSFRSYGASGKYDQEDSYSGKANEDYSMYSGYGGYGGYGAYAGNSMVNPAGFYGYGGGYGYGYGYGGQMFNMGYGAGGY

SHMGSGYGVAAAAAYGGGKSHGNGNNGSSSGKGNGTNGSGPDRYHPYQK*

>Athaliana_167_TAIR10.protein_primaryTranscriptOnly.fa_AT1G58470.1

MDYDRYKLFVGGIAKETSEEALKQYFSRYGAVLEAVVAKEKVTGKPRGFGFVRFANDCDVVKALRDTHFILGKPVDVRKA

IRKHELYQQPFSMQFLERKVQQMNGGLREMSSNGVTSRTKKIFVGGLSSNTTEEEFKSYFERFGRTTDVVVMHDGVTNRP

RGFGFVTYDSEDSVEVVMQSNFHELSDKRVEVKRAIPKEGIQSNNGNAVNIPPSYSSFQATPYVPEQNGYGMVLQFPPPV

FGYHHNVQAVQYPYGYQFTAQVANVSWNNPIMQPTGFYCAPPHPTPPPTNNLGYIQYMNGFDLSGTNISGYNPLAWPVTG

DAAGALIHQFVDLKLDVHSQAHQRMNGGNMGIPLQNGTYI*

>Athaliana_167_TAIR10.protein_primaryTranscriptOnly.fa_AT3G07810.2

MQSDNGKLFIGGISWDTNEERLKEYFSSFGEVIEAVILKDRTTGRARGFGFVVFADPAVAEIVITEKHNIDGRLVEAKKA

VPRDDQNMVNRSNSSSIQGSPGGPGRTRKIFVGGLPSSVTESDFKTYFEQFGTTTDVVVMYDHNTQRPRGFGFITYDSEE

AVEKVLLKTFHELNGKMVEVKRAVPKELSPGPSRSPLGAGYSYGVNRVNNLLNGYAQGFNPAAVGGYGLRMDGRFSPVGA

GRSGFANYSSGYGMNVNFDQGLPTGFTGGTNYNGNVDYGRGMSPYYIGNTNRFGPAVGYEGGNGGGNSSFFSSVTRNLWG

NNGGLNYNNNNTNSNSNTYMGGSSSGNNTLSGPFGNSGVNWGAPGGGNNAVSNENVKFGYGGNGESGFGLGTGGYAARNP

GANKAAPSSSFSSASATNNTGYDTAGLAEFYGNGAVYSDPTWRSPTPETEGPAPFSYGIGGGVPSSDVSARSSSPGYVGS

YSVNKRQPNRGEPSR*

>Athaliana_167_TAIR10.protein_primaryTranscriptOnly.fa_AT5G47620.4

MEMESCKLFIGGISWETSEDRLRDYFHSFGEVLEAVIMKDRATGRARGFGFVVFADPNVAERVVLLKHIIDGKILVDSIV

YNQLCRSDKCISLSEVVEAKKAVPRDDHVVFNKSNSSLQGSPGPSNSKKIFVGGLASSVTEAEFKKYFAQFGMITDVVVM

YDHRTQRPRGFGFISYDSEEAVDKVLQKTFHELNGKMVEVKLAVPKDMALNTMRNQMNVNSFGTSRISSLLNEYTQGFSP

SPISGYGVKPEVRYSPAVGNRGGFSPFGHGYGIELNFEPNQTQNYGSGSSGGFGRPFSPGYAASLGRFGSQMESGGASVG

NGSVLNAAPKNHLWGNGGLGYMSNSPISRSSFSGNSGMSSLGSIGDNWGTVARARSSYHGERGGVGLEAMRGVHVGGYSS

GSSILEADSLYSDSMWLSLPAKAEEGLGMGPLDFMSRGPAGYINRQPNGGIAA*

>Athaliana_167_TAIR10.protein_primaryTranscriptOnly.fa_AT5G55550.3

MESDLGKLFIGGISWDTDEERLRDYFSNYGDVVEAVIMRDRATGRARGFGFIVFADPCVSERVIMDKHIIDGRTVEAKKA

VPRDDQQVLKRHASPIHLMSPVHGGGGRTKKIFVGGLPSSITEEEFKNYFDQFGTIADVVVMYDHNTQRPRGFGFITFDS

DDAVDRVLHKTFHELNGKLVEVKRAVPKEISPVSNIRSPLASGVNYGGGSNRMPANSYFNNFAPGPGFYNSLGPVGRRFS

PVIGSGRNAVSAFGLGLNHDLSLNLNPSCDGTSSTFGYNRIPSNPYFNGASPNRYTSPIGHNRTESPYNSNNRDLWGNRT

DTAGPGWNLNVSNGNNRGNWGLPSSSAVSNDNNGFGRNYGTSSGLSSSPFNGFEGSIGELYRGGSVYSDSTWQQQQLPSQ

SSHELDNLSRAYGYDIDNVGSDPSANDPETYNGSYNVGNRQTNRGNKKIHLNKTCNLDTF*

>Creinhardtii_281_v5.5.protein_primaryTranscriptOnly.fa_Cre07.g330300.t1.2

MPDLCNMTDKQSAKVFVGGLSWETTGEKLRAYMENFGSVREAFVSYNRNNGRPRGFGFVVFESPEVADKVVATKHMIDRR

EVEAKRAVPKEDAPEEKQQGSAPQRTKKIFVGGLAPTVDEAQLRQHFSDFGTVEDAVVMYDHENKRPRGFGFVTFAEEEA

VERVFSHGAVQTIADKPIEVKSAVPRDQMPPTPRMQGSYYGQPPHRGGPPGPGYGPGPHRGPNFAGPPPYGPPGPGYPPP

YGGYGRPFNGRPPNNYGAYGPRNPGPGGPGGRGQPPMPGPLGPGAYGGYPQQRFPPANQGPGAQGGSGGNASAASGKVPP

MPNAYDVYSGQLNGVNNAAMLSSLYNIAGLQLPNGLPAAAAAAAAGVNPKQLNSLNNQLNALKLANFAANFANQPDGTYP

DDEAAAYAASQQEYAATAEAIQQAGLNGLAAVNADFNTLHDAGFTTAPAPGWSS*

>Creinhardtii_281_v5.5.protein_primaryTranscriptOnly.fa_Cre12.g560300.t1.2

MASEEQAPSSGTPGGKLFLGGLSWDTTEEKLKEYFAKYGEVQDVVVMRDRVTRKPRGFGFITFADPAAAQAACAEPHTID

GRQIDAKPSVPHGEGGQQPRSKKIFVGGLAPETDDAQLKAHFEQYGAVVEALVMVDHNSQRSRGFGFVTFAEEPSVEKVF

AAGQMHELGGKQVEVKSATPKGSGPQAGRGAGRGAVSAGGAYGGRGYGGGGGGGRGGYYQGQGYGGYGYGGFPGYGAGGF

GGYGAGMQGYGNYGMMGYGAGYGGMMGYGGYGGYGNYGYGNQQPYGNQPQGGRGGRGGAGGSAEGQY*

>Danio_rerio.GRCz11.pep.all.fa_ENSDARP00000117590.1

MNSNLAGDEIGKLFVGGLDWSTTQETLRNYFSQYGEVVDCVIMKDKSTNQSRGFGFVKFK

DPNCVRTVLDTKPHNLDGRNIDPKPCTPRGMQPEKTRTKDGWKGSKSDSNKSKKIFVGGI

PHNCGEAELRDYFNRFGVVTEVVMIYDAEKQRPRG

>Danio_rerio.GRCz11.pep.all.fa_ENSDARP00000130727.3

MSALPSTTLATEMNSNLAGDEIGKLFVGGLDWSTTQETLRNYFSQYGEVVDCVIMKDKST

NQSRGFGFVKFKDPNCVRTVLDTKPHNLDGRNIDPKPCTPRGMQPEKTRTKDGWKGSKSD

SNKSKKIFVGGIPHNCGEAELRDYFNRFGVVTEVVMIYDAEKQRPRGFGFITFEAEQSVD

QAVNMHFHDIMGKKVEVKKAEPRDSKAPAPGQLGANQWGPRAIISAANGWAAQPAPTWQQ

SYGPQGVWVSTGQPLAGYGPPLPAGRGAPAQAPSPFNAFLVAAPPTGGFGAPQGYPQQGF

STQPQFGYSFGTPQGDQFVAQGMPPPPATPGAGPLGFAPATTPSQDLSKAAPAQPDFSYS

QYGLGNYPQDPSAYGPTRPSHTYVQEEQGYSAGYGQDLSAFGHSFADPSQPTASYAAAPS

APSSGAPAAASSFGRGQNHNVQGFHPYRR

>Danio_rerio.GRCz11.pep.all.fa_ENSDARP00000111358.1

MNSNLAGDEIGKLFVGGLDWSTTQETLRNYFSQYGEVVDCVIMKDKSTNQSRGFGFVKFK

DPNCVRTVLDTKPHNLDGRNIDPKPCTPRGMQPEKTRTKDGWKGSKSDSNKSKKIFVGGI

PHNCGEAELRDYFNRFGVVTEVVMIYDAEKQRPRGFGFITFEAEQSVDQAVNMHFHDIMG

KKVEVKKAEPRDSKAPAPGQLGANQWGPRAIISAANGWAAQPAPTWQQSYGPQGVWVSTG

QPLGYGPPLPAGRGAPAQAPSPFNAFLVAAPPTGGFGAPQGYPQQGFSTQPQFGYSFGTP

QGDQFVAQGMPPPPATPGAGPLGFAPATTPSQDLSKAAPAQPDFSYSQYGYGQDLSAFGH

SFADPSQPTASYAAAPSAPSSGAP

>Danio_rerio.GRCz11.pep.all.fa_ENSDARP00000109996.1

MNSNLAGDEIGKLFVGGLDWSTTQETLRNYFSQYGEVVDCVIMKDKSTNQSRGFGFVKFK

DPNCVRTVLDTKPHNLDGRNIDPKPCTPRGMQPEKTRTKDGWKGSKSDSNKSKKIFVGGI

PHNCGEAELRDYFNRFGVVTEVVMIYDAEKQRPRGFGFITFEAEQSVDQAVNMHFHDIMG

KKVEVKKAEPRDSKAPAPGQLGANQWGPRAIISAANGWAAQPAPTWQQSYGPQGVWVSTG

QPLAGYGPPLPAGRGAPAQAPSPFNAFLVAAPPTGGFGAPQGYPQQGFSTQPQFGYSFGT

PQGDQFVAQGMPPPPATPGAGPLGFAPATTPSQDLSKAAPAQPDFSYSQYGYGQDLSAFG

HSFADPSQPTASYAAAPSAPSSGAP

>Danio_rerio.GRCz11.pep.all.fa_ENSDARP00000050648.8

MEGDGSQATSGSPNDSQHDPGKMFIGGLSWQTSPDSLRDYFSKFGEIRECMVMRDPTTKR

SRGFGFVTFADAASVDKVLAQPHHELDSKTIDPKVAFPRRAQPKMVTRTKKIFVGGLSAN

TVVEDVKQYFEQFGKVEDAMLMFDKTTNRHRGFGFVTFENEDVVEKVCEIHFHEINNKMV

ECKKAQPKEVMFPPGTRGRARGLPYTMDAFMLGMGMLSYPNIVATYGRGYTGFAPSYSYQ

FPGFPATAYGPVAAAAAVAAARGSGTNPPRPGAFPGASSPGPVADLYGPTSQDSGVGNYI

SAAS

>Danio_rerio.GRCz11.pep.all.fa_ENSDARP00000126843.2

MVMRDPTTKRSRGFGFVTFADAASVDKVLAQPHHELDSKTIDPKVAFPRRAQPKMVTRTK

KIFVGGLSANTVVEDVKQYFEQFGKVEDAMLMFDKTTNRHR

>Danio_rerio.GRCz11.pep.all.fa_ENSDARP00000126581.2

MVMRDPTTKRSRGFGFVTFADAASVDKVLAQPHHELDSKTIDPKVAFPRRAQPKMVTRTK

KIFVGGLSANTVVEDVKQYFEQFGKVEDAMLMFD

>Danio_rerio.GRCz11.pep.all.fa_ENSDARP00000126866.2

MNLKMFIGGLSWQTSPDSLRDYFSKFGEIRECMVMRDPTTKRSRGFGFVTFADAASVDKV

LAQPHHELDSKTIDPKVAFPRRAQPKMVTRTKKIFVGGLSANTVVEDVKQYFEQFGKVED

AMLMFDKTTNRHRGFGFVTFENEDVVEKVCEIHFHEINNKMVECKKAQPKEVMFPPGTRG

RARGLPYTMDAFMLGMGMLS

>Danio_rerio.GRCz11.pep.all.fa_ENSDARP00000123620.2

MVMRDPTTKRSRGFGFVTFADAASVDKVLAQPHHELDSKTIDPKVAFPRRAQPKMVTRTK

KIFVGGLSANTVVEDVKQYFEQFGKVEDAMLMFDKTTNRHRGFGFVTFENEDVVEKVCEI

HFHEINNKMVECKKAQPKEVMFPPGTRGRARGLPYTMDAFMLGMGMLSYPNIVATYGRGY

TGFAPSYSYQFP

>Danio_rerio.GRCz11.pep.all.fa_ENSDARP00000129590.1

XIFVGGLSANTVVEDVKQYFEQFGKVEDAMLMFDKTTNRHRGFGFVTFENEDVVEKVCEI

HFHEINNKMVECKKAQPKEVMFPPGTRGRARGLPYTMDAFMLGMGMLSYPNIVATYGRGY

TGFAPSYSYQFPGFPDYGFYSSPSDQRGPPFSFADYGSLGPQAAQLLQSEHATSACNSPL

QHLASPDQYKSPGTNPPRPGAFPGASSPGPVADLYGPTSQDSGVGNYISAASPQPGSGFG

HSIA

>Danio_rerio.GRCz11.pep.all.fa_ENSDARP00000090967.2

MEGDGSQATSGSPNDSQHDPGKMFIGGLSWQTSPDSLRDYFSKFGEIRECMVMRDPTTKR

SRGFGFVTFADAASVDKVLAQPHHELDSKTIDPKVAFPRRAQPKMVTRTKKIFVGGLSAN

TVVEDVKQYFEQFGKVEDAMLMFDKTTNRHRGFGFVTFENEDVVEKVCEIHFHEINNKMV

ECKKAQPKEVMFPPGTRGRARGLPYTMDAFMLGMGMLSYPNIVATYGRGYTGFAPSYSYQ

FPGFPATAYGPVAAAAAVAAARGSGRGARGRGGYAAYPQSTGPGFPDYGFYSSPSDQRGP

PFSFADYGSLGPQAAQLLQSEHATSACNSPLQHLASPDQYKSPGTNPPRPGAFPGASSPG

PVADLYGPTSQDSGVGNYISAASPQPGSGFGHSIAGPLIATAFTNGYH

>Danio_rerio.GRCz11.pep.all.fa_ENSDARP00000129111.1

XSKFGEIRECMVMRDPTTKRSRGFGFVTFADAASVDKVLAQPHHELDSKTIDPKVAFPRR

AQPKMVTRTKKIFVGGLSANTVVEDVKQYFEQFGKVEDAMLMFDKTTNRHRGFGFVTFEN

EDVVEKVCEIHFHEINNKMVECKKAQPKEVMFPPGTRGRARGLPYTMDAFMLGMGMLSYP

NIVATYGRGYTGFAPSYSYQFPEFIRRPGEDWPEFLIGSVYR

>Danio_rerio.GRCz11.pep.all.fa_ENSDARP00000128256.1

XVECKKAQPKEVMFPPGTRGRARGLPYTMDAFMLGMGMLSYPNIVATYGRGYTGFAPSYS

YQFPGFPATAYGPVAAAAAVAAARGSGFPDYGFYSSPSDQRGPPFSFADYGSLGPQAAQL

LQSEHATSACNSPLQHLASPDQYKSPGTNPPRPGAFPGASSPGPVADLYGPTSQDSGVGN

YISAASPQPGSGFGHSIAGPLIATAFTNGYH

>Danio_rerio.GRCz11.pep.all.fa_ENSDARP00000127772.1

XGLSANTVVEDVKQYFEQFGKVEDAMLMFDKTTNRHRGFGFVTFENEDVVEKVCEIHFHE

INNKMVECKKAQPKEVMFPPGTRGRARGLPYTMDAFMLGMGMLSYPNIVATYGRGYTGFA

PSYSYQFPGTNPPRPGAFPGASSPGPVADLYGPTSQDSGVGNYISAASP

>Danio_rerio.GRCz11.pep.all.fa_ENSDARP00000121659.1

IDPKVAFPRRAQPKLVTRTKKIFVGGLSVNTTIEDVKQYFDQFGKVDDAMLMFDKTTNRH

RGFGFVTFENEDVVEKVCEIHFHEINNKMVECKKAQPKEVMSPTGSSRGRARVMPYGMDA

FMLGIGMLGYPGFQATTYAGRSYTSLTPGYTYQFPEFHLERTPLLTSPHPPELTAIPLTA

YNPMAAAAAAAAVVRGSTPSRSAGFLGTSSPGPM

>Danio_rerio.GRCz11.pep.all.fa_ENSDARP00000088198.4

XGLKDYFCKFGEVKESMVMRDPVTKRSRGFGFVTFVDQAGVDKVLAQTRHELDSKTIDPK

VAFPRRAQPKLVTRTKKIFVGGLSVNTTIEDVKQYFDQFGKVDDAMLMFDKTTNRHRGFG

FVTFENEDVVEKVCEIHFHEINNKMVECKKAQPKEVMSPTGSSRGRARVMPYGMDAFMLG

IGMLGYPGFQATTYAGRSYTSLTPGYTYQFPAIPLTAYNPMAAAAAAAAVVRGSTPSRSA

GFLGTSSPGPMADLYGTANQESAVSSYLSAASPAPSTGFSHSLGGPLIATAFTNGYH

>Danio_rerio.GRCz11.pep.all.fa_ENSDARP00000090247.2

MEADASQVTSGSLNDSQHDPGKMFIGGLSWQTSPDSLRDYFCKFGEIRECMVMRDPTTKR

SRGFGFITFADVSSVDKVLAQPHHELDSKTIDPKVAFPRRAQPKMVTRTKKIFVGGLSAS

TVVEDVKQYFEQFGKVEDAMLMFDKTTNRHRGFGFVTFENEDIVEKVCEIHFHEINNKMV

ECKKAQPKEVMFPPGTRGRARSLPYTMDAFMLGMGMLSYPNIVATYGRGYTGFSPSYSYQ

FPGFPATAYGPVAAAAVAAARGSGFPDYGFYSGPGDQRAAPCSFADYATLGPHTGQMLQS

EHVTSTCNSPSQHHPSPDHFKSSGANPPRPGGFPGANSPGPVADLYGPSSQDSAVGNYIS

AASPQPGSGFNHSIAGPLIATAFTNGYH

>Danio_rerio.GRCz11.pep.all.fa_ENSDARP00000123512.1

MEADASQVTSGSLNDSQHDPGKMFIGGLSWQTSPDSLRDYFCKFGEIRECMVMRDPTTKR

SRGFGFITFADVSSVDKVLAQPHHELDSKTIDPKVAFPRRAQPKMVTRTKKIFVGGLSAS

TVVEDVKQYFEQFGKVEDAMLMFDKTTNRHRGFGFVTFENEDIVEKVCEIHFHEINNKMV

ECKKAQPKEVMFPPGTRGRARSLPYTMDAFMLGMGMLSYPNIVATYGRGYTGFSPSYSYQ

FPGFPATAYGPVAAAAVAAARGSGFPDYGFYSGPGDQRAAPCSFADYATLGPHTGQMLQS

EHVTSTCNSPSQHHPSPDHFKSSGANPPRPGGFPGANSPGPVADLYGPSSQDSAVGNYIS

AASPQPGSGFNHSIAGPLIATAFTNGYH

>Danio_rerio.GRCz11.pep.all.fa_ENSDARP00000146314.1

FGKQSLNGSTLRNYFSQYGEVVDCVIMKDKSTNQSRGFGFVKFKDPNCVRTVLDTKPHNL

DGRNIDPKPCTPRGMQPEKTRTKDGWKGSKSDSNKSKKIFVGGIPHNCGEAELRDYFNRF

GVVTEVVMIYDAEKQRPRGFGFITFEAEQSVDQAVNMHFHDIMGKKVEVKKAEPRDSKAP

APGQLGANQWGPRAIISAANGWAAQPAPTWQQSYGPQGVWVSTGQPLAGYGPPLPAGRGA

PAQAPSPFNAFLVAAPPTGGFGAPQGYPQQGFSTQPQFGYSFGTPQGDQFVAQGMPPPPA

TPGAGPLGFAPATTPSQDLSKAAPAQPDFSYSQYGLGNYPQDPSAYGPTRPSHTYVQEEQ

GYSAGYGQDLSAFGHSFADPSQPTASYAAAPSAPSSGAPAAASSFGRGQNHNVQGFHPYR

R

>Danio_rerio.GRCz11.pep.all.fa_ENSDARP00000151028.1

MSPGLCVLKRVWDCIDIIDWSSQTLRNYFSQYGEVVDCVIMKDKSTNQSRGFGFVKFKDP

NCVRTVLDTKPHNLDGRNIDPKPCTPRGMQPEKTRTKDGWKGSKSDSNKSKKIFVGGIPH

NCGEAELRDYFNRFGVVTEVVMIYDAEKQRPRGFGFITFEAEQSVDQAVNMHFHDIMGKK

VEVKKAEPRDSKAPAPGQLGANQWGPRAIISAANGWAAQPAPTWQQSYGPQGVWVSTGQP

LAGYGPPLPAGRGAPAQAPSPFNAFLVAAPPTGGFGAPQGYPQQGFSTQPQFGYSFGTPQ

GDQFVAQGMPPPPATPGAGPLGFAPATTPSQDLSKAAPAQPDFSYSQYGYGQDLSAFGHS

FADPSQPTASYAAAPSAPSSGAPAAASSFGRGQNHNVQGFHPYRR

>Danio_rerio.GRCz11.pep.all.fa_ENSDARP00000146821.1

TFENEDIVEKVCEIHFHEINNKMVECKKAQPKEVMFPPGTRGRARSLPYTMDAFMLGMGM

LSYPNIVATYGRGYTGFSPSYSYQFPGFPATAYGPVAAAAVAAARGSGRNAGPGKHSLPP

CVCVSYTGANPPRPGGFPGANSPGPVADLYGPSSQDSAVGNYISAASPQPGSGFNHSIAP

IIESLFTCVDVHLFSFESNFSNITD

>Danio_rerio.GRCz11.pep.all.fa_ENSDARP00000156045.1

MEADASQVTSGSLNDSQHDPGKMFIGGLSWQTSPDSLRDYFCKFGEIRECMVMRDPTTKR

SRGFGFITFADVSSVDKVLAQPHHELDSKTIDPKVAFPRRAQPKMVTRTKKIFVGGLSAS

TVVEDVKQYFEQFGKVRTRTLDATTNRIKLIIKSFGFVTFENEDIVEKVCEIHFHEINNK

MVECKKAQPKEVMFPPGTRGRARSLPYTMDAFMLGMGMLSYPNIVATYGRGYTGFSPSYS

YQFPGEHLSLSIYLSIYLSIYLSIYLSIYLSIYLSIYLSVCLSVCLSVCLSVCLSVCLSV

CLSVCLSVCLSVCLSVELIYHKLIDLPFYLSFQEFVHVSCYICA

>Danio_rerio.GRCz11.pep.all.fa_ENSDARP00000135610.1

MAYRVSSLSFVCVCVCVCVCVCFCLQVDDAMLMFDKTTNRHRGFGFVTFENEDVVEKVCE

IHFHEINNKMVECKKAQPKEVMSPTGTARGRSRVMPYGMDAFMLGIGMLGYPGFQAATYA

SRGYTGITPGYTYQFPEFHLERTPLLTSTHPPEITAIPLTAYGPMAAAAAAAVVRGSTPS

RFLGTSSPGPMADLYAAASQESAVSSYISAASPAPSTGFGHGLGGPLIATAFTNGYH

>Drosophila_melanogaster.BDGP6.pep.all.fa_FBpp0301711

MVTRTKKIFVGGLSAPTTLEDVKSYFEQFGPIEDAMLMFDKQTNRHRGFGFVTFQSEDVV

DKVCEIHFHEINNKMVECKKAQPKEVMLPANLAKTRAAGRSAYDNIMWGLGTLPEGFPAA

AYAAYAAGRGYSGYPSFGLPYPTVDLTNGQLTPNNNTPLLTQALSVMNNYQAAAAAAHSY

GPSASPHAAAANTATRQGFPSTNSPGPTIDMYSSAAADNVGYVQATSPQPSGFPIAVSRA

PLNYSPGPLIPAAFTNGYH

>Drosophila_melanogaster.BDGP6.pep.all.fa_FBpp0301712

MVTRTKKIFVGGLSAPTTLEDVKSYFEQFGPIEDAMLMFDKQTNRHRGFGFVTFQSEDVV

DKVCEIHFHEINNKMVECKKAQPKEVMLPANLAKTRAAGRSAYDNIMWGLGTLPEGFPAA

AYAAYAAGRGYSGYPSFGLPYPTGLTICGIGRSYGTACSATPGRSRGATAGAASGVAATG

VAPSGGAPSHAQSAYHQQALTLPLATALTARSAAIKSQFLGNINF

>Drosophila_melanogaster.BDGP6.pep.all.fa_FBpp0075045

MDIKIEQQQQQQQVELGPCSPSEVPNDPGKMFIGGLSWQTSPESLRDYFGRYGDISEAMV

MKDPTTRRSRGFGFVTFSDPNSVDKVLTQGTHELDGKKVDPKVAFPRRAHPKMVTRTKKI

FVGGLSAPTTLEDVKSYFEQFGPIEDAMLMFDKQTNRHRGFGFVTFQSEDVVDKVCEIHF

HEINNKMVECKKAQPKEVMLPANLAKTRAAGRSAYDNIMWGLGTLPEGFPAAAYAAYAAG

RGYSGYPSFGLPYPTVDLTNGQLTPNNNTPLLTQALSVMNNYQAAAAAAHSYGPSASPHA

AAANTATRQGFPSTNSPGPTIDMYSSAAADNVGYVQATSPQPSGFPIAVSRAPLNYSPIY

SLNFAMANI

>Drosophila_melanogaster.BDGP6.pep.all.fa_FBpp0305994

MDIKIEQQQQQQQVELGPCSPSEVPNDPGKMFIGGLSWQTSPESLRDYFGRYGDISEAMV

MKDPTTRRSRGFGFVTFSDPNSVDKVLTQGTHELDGKKVDPKVAFPRRAHPKMVTRTKKI

FVGGLSAPTTLEDVKSYFEQFGPIEDAMLMFDKQTNRHRGFGFVTFQSEDVVDKVCEIHF

HEINNKMVECKKAQPKEVMLPANLAKTRAAGRSAYDNIMWGLGTLPEGFPAAAYAAYAAG

RGYSGYPSFGLPYPTVDLTNGQLTPNNNTPLLTQALSVMNNYQAAAAAAHSYGPSASPHA

AAANTATRQGFPSTNSPGPTIDMYSSAAADNVGYVQATSPQPSGFPIAVSRAPLNYSPGP

LIPAAFTNGYH

>Drosophila_melanogaster.BDGP6.pep.all.fa_FBpp0112339

MDIKIEQQQQQQQVELGPCSPSEVPNDPGKMFIGGLSWQTSPESLRDYFGRYGDISEAMV

MKDPTTRRSRGFGFVTFSDPNSVDKVLTQGTHELDGKKVDPKVAFPRRAHPKMVTRTKKI

FVGGLSAPTTLEDVKSYFEQFGPIEDAMLMFDKQTNRHRGFGFVTFQSEDVVDKVCEIHF

HEINNKMVECKKAQPKEVMLPANLAKTRAAGRSAYGELVVWGSSHAHSTAATSAAAAGLL

PSSLAAAASVLQQQQQQQQQQQQQQQQQQQHHQQLHHTHPQSPAHSHAHHPHTHSHSQLL

GSLRYTPYPLPAHLSAAAVVVAQQQHHHQQQQQQQQQQQQQQVVAAQQQHQQQQSLAQVV

AAAAGAAPGLLPLANPAPPATPSLLQFAAGQSNAALANSLYADAAAVVGYKRLLAAAAVS

SGLRAPTTALGALQAAAANVPAAAAAAQLQQAQLRQNAALAAAHYPLSELLAMQGGMEMG

AGANSAAAAAASLYQLPGI

>Drosophila_melanogaster.BDGP6.pep.all.fa_FBpp0112340

MVTRTKKIFVGGLSAPTTLEDVKSYFEQFGPIEDAMLMFDKQTNRHRGFGFVTFQSEDVV

DKVCEIHFHEINNKMVECKKAQPKEVMLPANLAKTRAAGRSAYDNIMWGLGTLPEGFPAA

AYAAYAAGRGYSGYPSFGLPYPTVDLTNGQLTPNNNTPLLTQALSVMNNYQAAAAAAHSY

GPSASPHAAAANTATRQGFPSTNSPGPTIDMYSSAAADNVGYVQATSPQPSGFPIAVSRA

PLNYSPIYSLNFAMANI

>Drosophila_melanogaster.BDGP6.pep.all.fa_FBpp0288505

MVTRTKKIFVGGLSAPTTLEDVKSYFEQFGPIEDAMLMFDKQTNRHRGFGFVTFQSEDVV

DKVCEIHFHEINNKMVECKKAQPKEVMLPANLAKTRAAGRSAYDNIMWGLGTLPEGFPAA

AYAAYAAGRGYSGYPSFGLPYPTASVMGVVPENRILIGDYMA

>Drosophila_melanogaster.BDGP6.pep.all.fa_FBpp0078975

MEEDERGKLFVGGLSWETTQENLSRYFCRFGDIIDCVVMKNNESGRSRGFGFVTFADPTN

VNHVLQNGPHTLDGRTIDPKPCNPRTLQKPKKGGGYKVFLGGLPSNVTETDLRTFFNRYG

KVTEVVIMYDQEKKKSRGFGFLSFEEESSVEHVTNERYINLNGKQVEIKKAEPRDGSGGQ

NSNNSTVGGAYGKLGNECSHWGPHHAPINMMQGQNGQMGGPPLNMPIGAPNMMPGYQGWG

TSPQQQQYGYGNSGPGSYQGWGAPPGPQGPPPQWSNYAGPQQTQGYGGYDMYNSTSTGAP

SGPSGGGSWNSWNMPPNSAGPTGAPGAGAGTATDMYSRAQAWATGGPSTTGPVGGMPRTG

PGNSASKSGSEYDYGGYGSGYDYDYSNYVKQEGASNYGAGPRSAYGNDSSTQPPYATSQA

V

>Drosophila_melanogaster.BDGP6.pep.all.fa_FBpp0078974

MEEDERGKLFVGGLSWETTQENLSRYFCRFGDIIDCVVMKNNESGRSRGFGFVTFADPTN

VNHVLQNGPHTLDGRTIDPKPCNPRTLQKPKKGGGYKVFLGGLPSNVTETDLRTFFNRYG

KVTEVVIMYDQEKKKSRGFGFLSFEEESSVEHVTNERYINLNGKQVEIKKAEPRDGSGGQ

NSNNSTVGGAYGKLGNECSHWGPHHAPINMMQGQNGQMGGPPLNMPIGAPNMMPGYQGWG

TSPQQQQYGYGNSGPGSYQGWGAPPGPQGPPPQWSNYAGPQQTQGYGGYDMYNSTSTGAP

SGPSGGGSWNSWNMPPNSAGPTGAPGAGAGTATDMYSRAQAWATGGPSTTGPVGGMPRTG

PGNSASKSGSEYDYGGYGSGYDYDYSNYVKQEGASNYGAGPRSAYGNDSSTQPPYATSQA

V

>Drosophila_melanogaster.BDGP6.pep.all.fa_FBpp0297874

MEEDERGKLFVGGLSWETTQENLSRYFCRFGDIIDCVVMKNNESGRSRGFGFVTFADPTN

VNHVLQNGPHTLDGRTIDPKPCNPRTLQKPKKGGGYKVFLGGLPSNVTETDLRTFFNRYG

KVTEVVIMYDQEKKKSRGFGFLSFEEESSVEHVTNERYINLNGKQVEIKKAEPRDGSGGQ

NSNNSTVGGAYGKLGNECSHWGPHHAPINMMQGQNGQMGGPPLNMPIGAPNMMPGYQGWG

TSPQQQQYGYGNSGPGSYQGWGAPPGPQGPPPQWSNYAGPQQTQGYGGYDMYNSTSTGAP

SGPSGGGSWNSWNMPPNSAGPTGAPGAGAGTATDMYSRAQAWATGGPSTTGPVGGMPRTG

PGNSASKSGSEYDYGGYGSGYDYDYSNYVKQEGASNYGAGPRSAYGNDSSTQPPYATSQA

V

>Drosophila_melanogaster.BDGP6.pep.all.fa_FBpp0297875

MEEDERGKLFVGGLSWETTQENLSRYFCRFGDIIDCVVMKNNESGRSRGFGFVTFADPTN

VNHVLQNGPHTLDGRTIDPKPCNPRTLQKPKKGGGYKVFLGGLPSNVTETDLRTFFNRYG

KVTEVVIMYDQEKKKSRGFGFLSFEEESSVEHVTNERYINLNGKQVEIKKAEPRDGSGGQ

NSNNSTVGGAYGKLGNECSHWGPHHAPINMMQGQNGQMGGPPLNMPIGAPNMMPGYQGWG

TSPQQQQYGYGNSGPGSYQGWGAPPGPQGPPPQWSNYAGPQQTQGYGGYDMYNSTSTGAP

SGPSGGGSWNSWNMPPNSAGPTGAPGAGAGTATDMYSRAQAWATGGPSTTGPVGGMPRTG

PGNSASKSGSEYDYGGYGSGYDYDYSNYVKQEGASNYGAGPRSAYGNDSSTQPPYATSQA

V

>Drosophila_melanogaster.BDGP6.pep.all.fa_FBpp0290617

MEEDERGKLFVGGLSWETTQENLSRYFCRFGDIIDCVVMKNNESGRSRGFGFVTFADPTN

VNHVLQNGPHTLDGRTIDPKPCNPRTLQKPKKGGGYKVFLGGLPSNVTETDLRTFFNRYG

KVTEVVIMYDQEKKKSRGFGFLSFEEESSVEHVTNERYINLNGKQVEIKKAEPRDGSGGQ

NSNNSTVGGAYGKLGNECSHWGPHHAPINMMQGQNGQMGGPPLNMPIGAPNMMPGYQGWG

TSPQQQQYGYGNSGPGSYQGWGAPPGPQGPPPQWSNYAGPQQTQGYGGYDMYNSTSTGAP

SGPSGGGSWNSWNMPPNSAGPTGAPGAGAGTATDMYSRAQAWATGGPSTTGPVGGMPRTG

PGNSASKSGSEYDYGGYGSGYDYDYSNYVKQEGASNYGAGPRSAYGNDSSTQPPYATSQA

V

>Drosophila_melanogaster.BDGP6.pep.all.fa_FBpp0078976

MEEDERGKLFVGGLSWETTQENLSRYFCRFGDIIDCVVMKNNESGRSRGFGFVTFADPTN

VNHVLQNGPHTLDGRTIDPKPCNPRTLQKPKKGGGYKVFLGGLPSNVTETDLRTFFNRYG

KVTEVVIMYDQEKKKSRGFGFLSFEEESSVEHVTNERYINLNGKQVEIKKAEPRDGSGGQ

NSNNSTVGGAYGKLGNECSHWGPHHAPINMMQGQNGQMGGPPLNMPIGAPNMMPGYQGWG

TSPQQQQYGYGNSGPGSYQGWGAPPGPQGPPPQWSNYAGPQQTQGYGGYDMYNSTSTGAP

SGPSGGGSWNSWNMPPNSAGPTGAPGAGAGTATDMYSRAQAWATGGPSTTGPVGGMPRTG

PGNSASKSGSEYDYGGYGSGYDYDYSNYVKQEGASNYGAGPRSAYGNDSSTQPPYATSQA

V

>Drosophila_melanogaster.BDGP6.pep.all.fa_FBpp0297873

MEEDERGKLFVGGLSWETTQENLSRYFCRFGDIIDCVVMKNNESGRSRGFGFVTFADPTN

VNHVLQNGPHTLDGRTIDPKPCNPRTLQKPKKGGGYKVFLGGLPSNVTETDLRTFFNRYG

KVTEVVIMYDQEKKKSRGFGFLSFEEESSVEHVTNERYINLNGKQVEIKKAEPRDGSGGQ

NSNNSTVGGAYGKLGNECSHWGPHHAPINMMQGQNGQMGGPPLNMPIGAPNMMPGYQGWG

TSPQQQQYGYGNSGPGSYQGWGAPPGPQGPPPQWSNYAGPQQTQGYGGYDMYNSTSTGAP

SGPSGGGSWNSWNMPPNSAGPTGAPGAGAGTATDMYSRAQAWATGGPSTTGPVGGMPRTG

PGNSASKSGSEYDYGGYGSGYDYDYSNYVKQEGASNYGAGPRSAYGNDSSTQPPYATSQA

V

>Drosophila_melanogaster.BDGP6.pep.all.fa_FBpp0290618

MEEDERGKLFVGGLSWETTQENLSRYFCRFGDIIDCVVMKNNESGRSRGFGFVTFADPTN

VNHVLQNGPHTLDGRTIDPKPCNPRTLQKPKKGGGYKVFLGGLPSNVTETDLRTFFNRYG

KVTEVVIMYDQEKKKSRGFGFLSFEEESSVEHVTNERYINLNGKQVEIKKAEPRDGSGGQ

NSNNSTVGGAYGKLGNECSHWGPHHAPINMMQGQNGQMGGPPLNMPIGAPNMMPGYQGWG

TSPQQQQYGYGNSGPGSYQGWGAPPGPQGPPPQWSNYAGPQQTQGYGGYDMYNSTSTGAP

SGPSGGGSWNSWNMPPNSAGPTGAPGAGAGTATDMYSRAQAWATGGPSTTGPVGGMPRTG

PGNSASKSGSEYDYGGYGSGYDYDYSNYVKQEGASNYGAGPRSAYGNDSSTQPPYATSQA

V

>Drosophila_melanogaster.BDGP6.pep.all.fa_FBpp0305821

MENAAAAAAAAAAGLIDPHHNRDLHQALVASIANNSVAAIGGGLTTAAVLKSAAQQSQQA

VQQNQNAVVVTPGLEQPKQEPAQQAALALLKENVNASAGAGQNNGQAAMGGSNKSGSSGR

STPSLSGGSGSDPAPGKLFVGGLSWQTSSDKLKEYFNMFGTVTDVLIMKDPVTQRSRGFG

FITFQEPCTVEKVLKVPIHTLDGKKIDPKHATPKNRPRQANKTKKIFVGGVSQDTSAEEV

KAYFSQFGPVEETVMLMDQQTKRHRGFGFVTFENEDVVDRVCEIHFHTIKNKKVECKKAQ

PKEAVTPAAQLLQKRIMLGTLGVQLPTAPGQLIGARGAGVATMNPLAMLQNPTQLLQSPA

AAAAAQQAALISQNPFQVQNAAAAASIANQAGFGKLLTTYPQTALHSVRYAPYSIPASAA

TANAALMQAHQAQSVAAAAHHHQQQQQQQHHHQQQTHNAHVAAAQQQQQSHHNAVSNPAS

QAHSAAAAAALAANAANGAGAAGAHSLAAAAQQAGLMAGNPLNAAAAAAAAAANPAAAYS

NYALANVDMSSFQGVDWSTMYGMGMYV

>Drosophila_melanogaster.BDGP6.pep.all.fa_FBpp0084256

MHALQEGATVLHHQQPPPPTSGEDHLLTADSFFYARSNPMENAAAAAAAAAAGLIDPHHN

RDLHQALVASIANNSVAAIGGGLTTAAVLKSAAQQSQQAVQQNQNAVVVTPGLEQPKQEP

AQQAALALLKENVNASAGAGQNNGQAAMGGSNKSGSSGRSTPSLSGGSGSDPAPGKLFVG

GLSWQTSSDKLKEYFNMFGTVTDVLIMKDPVTQRSRGFGFITFQEPCTVEKVLKVPIHTL

DGKKIDPKHATPKNRPRQANKTKKIFVGGVSQDTSAEEVKAYFSQFGPVEETVMLMDQQT

KRHRGFGFVTFENEDVVDRVCEIHFHTIKNKKVECKKAQPKEAVTPAAQLLQKRIMLGTL

GVQLPTAPGQLIGARGAGVATMNPLAMLQNPTQLLQSPAAAAAAQQAALISQNPFQVQNA

AAAASIANQAGFGKLLTTYPQTALHSVRYAPYSIPASAATANAALMQAHQAQSVAAAAHH

HQQQQQQQHHHQQQTHNAHVAAAQQQQQSHHNAVSNPASQAHSAAAAAALAANAANGAGA

AGAHSLAAAAQQAGLMAGNPLNAAAAAAAAAANPAAAYSNYALANVDMSSFQGVDWSTMY

GMGMYV

>Drosophila_melanogaster.BDGP6.pep.all.fa_FBpp0293749

MLFENPAVAAKLPFPYNVPPPLQAAAAAAAAVPNLSNPMENAAAAAAAAAAGLIDPHHNR

DLHQALVASIANNSVAAIGGGLTTAAVLKSAAQQSQQAVQQNQNAVVVTPGLEQPKQEPA

QQAALALLKENVNASAGAGQNNGQAAMGGSNKSGSSGRSTPSLSGGSGSDPAPGKLFVGG

LSWQTSSDKLKEYFNMFGTVTDVLIMKDPVTQRSRGFGFITFQEPCTVEKVLKVPIHTLD

GKKIDPKHATPKNRPRQANKTKKIFVGGVSQDTSAEEVKAYFSQFGPVEETVMLMDQQTK

RHRGFGFVTFENEDVVDRVCEIHFHTIKNKKVECKKAQPKEAVTPAAQLLQKRIMLGTLG

VQLPTAPGQLIGARGAGVATMNPLAMLQNPTQLLQSPAAAAAAQQAALISQNPFQVQNAA

AAASIANQAGFGKLLTTYPQTALHSVRYAPYSIPASAATANAALMQAHQAQSVAAAAHHH

QQQQQQQHHHQQQTHNAHVAAAQQQQQSHHNAVSNPASQAHSAAAAAALAANAANGAGAA

GAHSLAAAAQQAGLMAGNPLNAAAAAAAAAANPAAAYSNYALANVDMSSFQGVDWSTMYG

MGMYV

>Drosophila_melanogaster.BDGP6.pep.all.fa_FBpp0305819

MLFENPAVAAKLPFPYNVPPPLQAAAAAAAAVPNLRSVSEMNATSLYAGNPMENAAAAAA

AAAAGLIDPHHNRDLHQALVASIANNSVAAIGGGLTTAAVLKSAAQQSQQAVQQNQNAVV

VTPGLEQPKQEPAQQAALALLKENVNASAGAGQNNGQAAMGGSNKSGSSGRSTPSLSGGS

GSDPAPGKLFVGGLSWQTSSDKLKEYFNMFGTVTDVLIMKDPVTQRSRGFGFITFQEPCT

VEKVLKVPIHTLDGKKIDPKHATPKNRPRQANKTKKIFVGGVSQDTSAEEVKAYFSQFGP

VEETVMLMDQQTKRHRGFGFVTFENEDVVDRVCEIHFHTIKNKKVECKKAQPKEAVTPAA

QLLQKRIMLGTLGVQLPTAPGQLIGARGAGVATMNPLAMLQNPTQLLQSPAAAAAAQQAA

LISQNPFQVQNAAAAASIANQAGFGKLLTTYPQTALHSVRYAPYSIPASAATANAALMQA

HQAQSVAAAAHHHQQQQQQQHHHQQQTHNAHVAAAQQQQQSHHNAVSNPASQAHSAAAAA

ALAANAANGAGAAGAHSLAAAAQQAGLMAGNPLNAAAAAAAAAANPAAAYSNYALANVDM

SSFQGVDWSTMYGMGMYV

>Homo_sapiens.GRCh38.pep.all.fa_ENSP00000337132.5

MNNSGADEIGKLFVGGLDWSTTQETLRSYFSQYGEVVDCVIMKDKTTNQSRGFGFVKFKD

PNCVGTVLASRPHTLDGRNIDPKPCTPRGMQPERTRPKEGWQKGPRSDNSKSNKIFVGGI

PHNCGETELREYFKKFGVVTEVVMIYDAEKQRPRGFGFITFEDEQSVDQAVNMHFHDIMG

KKVEVKRAEPRDSKSQAPGQPGASQWGSRVVPNAANGWAGQPPPTWQQGYGPQGMWVPAG

QAIGGYGPPPAGRGAPPPPPPFTSYIVSTPPGGFPPPQGFPQGYGAPPQFSFGYGPPPPP

PDQFAPPGVPPPPATPGAAPLAFPPPPSQAAPDMSKPPTAQPDFPYGQYGLGSYSPAPPG

CGPHFVYSLMVRLSSDVA

>Homo_sapiens.GRCh38.pep.all.fa_ENSP00000233078.3

MNNSGADEIGKLFVGGLDWSTTQETLRSYFSQYGEVVDCVIMKDKTTNQSRGFGFVKFKD

PNCVGTVLASRPHTLDGRNIDPKPCTPRGMQPERTRPKEGWQKGPRSDNSKSNKIFVGGI

PHNCGETELREYFKKFGVVTEVVMIYDAEKQRPRGFGFITFEDEQSVDQAVNMHFHDIMG

KKVEVKRAEPRDSKSQAPGQPGASQWGSRVVPNAANGWAGQPPPTWQQGYGPQGMWVPAG

QAIGGYGPPPAGRGAPPPPPPFTSYIVSTPPGGFPPPQGFPQGYGAPPQFSFGYGPPPPP

PDQFAPPGVPPPPATPGAAPLAFPPPPSQAAPDMSKPPTAQPDFPYGQYAGYGQDLSGFG

QGFSDPSQQPPSYGGPSVPGSGGPPAGGSGFGRGQNHNVQGFHPYRR

>Homo_sapiens.GRCh38.pep.all.fa_ENSP00000467680.2

MNNSGADEIGKLFVGGLDWSTTQETLRSYFSQYGEVVDCVIMKDKTTNQSRGFGFVKFKD

PNCVGTVLASRPHTLDGRNIDPKPCTPRGMQPERTRPKEGWQKGPRSDNSKSNKIFVGGI

PHNCGETELREYFKKFGVVTEVVMIYDAEKQRPRGFGFITFEDEQSVDQAVNMHFHDIMG

KKVEVKRAEPRDSKSQAPGQPGASQWGSRVVPNAANGWAGQPPPTWQQGYGPQGMWVPAG

QAIGGYGPPPAGRGAPPPPPPFTSYIVSTPPGGFPPPQGFPQGYGAPPQFSFGYGPPPPP

PDQFAPPGVPPPPATPGAAPLAFPPPPSQAAPDMSKPPTAQPDFPYGQYGYGQDLSGFGQ

GFSDPSQQPPSYGGPSVPGSGGPPAGGSGFGRGQNHNVQGFHPYRR

>Homo_sapiens.GRCh38.pep.all.fa_ENSP00000465433.2

MNNSGADEIGKLFVGGLDWSTTQETLRSYFSQYGEVVDCVIMKDKTTNQSRGFGFVKFKD

PNCVGTVLASRPHTLDGRNIDPKPCTPRGMQPERTRPKEGWKGPRSDNSKSNKIFVGGIP

HNCGETELREYFKKFGVVTEVVMIYDAEKQRPRGFGFITFEDEQSVDQAVNMHFHDIMGK

KVEVKRAEPRDSKSQAPGQPGASQWGSRVVPNAANGWAGQPPPTWQQGYGPQGMWVPAGQ

AIGGYGPPPAGRGAPPPPPPFTSYIVSTPPGGFPPPQGFPQGYGAPPQFSFGYGPPPPPP

DQFAPPGVPPPPATPGAAPLAFPPPPSQAAPDMSKPPTAQPDFPYGQYAGYGQDLSGFGQ

GFSDPSQQPPSYGGPSVPGSGGPPAGGSGFGRGQNHNVQGFHPYRR

>Homo_sapiens.GRCh38.pep.all.fa_ENSP00000467613.2

MNNSGADEIGKLFVGGLDWSTTQETLRSYFSQYGEVVDCVIMKDKTTNQSRGFGFVKFKD

PNCVGTVLASRPHTLDGRNQKGPRSDNSKSNKIFVGGIPHNCGETELREYFKKFGVVTEV

VMIYDAEKQRPRGFGFITFEDEQSVDQAVNMHFHDIMGKKVEVKRAEPRDSKSQAPGQPG

ASQWGSRVVPNAANGWAGQPPPTWQQGYGPQGMWVPAGQAIGGYGPPPAGRGAPPPPPPF

TSYIVSTPPGGFPPPQGFPQGYGAPPQFSFGYGPPPPPPDQFAPPGVPPPPATPGAAPLA

FPPPPSQAAPDMSKPPTAQPDFPYGQY

>Homo_sapiens.GRCh38.pep.all.fa_ENSP00000257552.2

METDAPQPGLASPDSPHDPCKMFIGGLSWQTTQEGLREYFGQFGEVKECLVMRDPLTKRS

RGFGFVTFMDQAGVDKVLAQSRHELDSKTIDPKVAFPRRAQPKMVTRTKKIFVGGLSVNT

TVEDVKQYFEQFGKVDDAMLMFDKTTNRHRGFGFVTFESEDIVEKVCEIHFHEINNKMVE

CKKAQPKEVMSPTGSARGRSRVMPYGMDAFMLGIGMLGYPGFQATTYASRSYTGLAPGYT

YQFPEFRVERTPLPSAPVLPELTAIPLTAYGPMAAAAAAAAVVRGTGSHPWTMAPPPGST

PSRTGGFLGTTSPGPMAELYGAANQDSGVSSYISAASPAPSTGFGHSLGGPLIATAFTNG

YH

>Homo_sapiens.GRCh38.pep.all.fa_ENSP00000446710.1

XDQAGVDKVLAQSRHELDSKTIDPKVAFPRRAQPKMVTRTKKIFVGGLSVNTTVEDVKQY

FEQFGKVDDAMLMFDKTTNRHRGFGFVTFESEDIVEKVCEIHFHEINNKMVECKKAQPKE

VMSPTGSARGRSRVMPYGMDAFMLGIGMLGYPGFQATTYASRSYTGLAPGYTYQFPAIPL

TAYGPMAAAAAAAAVVRGTGSHPWTMAPPPGSTPSRTGGFLGTTSPGPMAELY

>Homo_sapiens.GRCh38.pep.all.fa_ENSP00000414671.2

MFIGGLSWQTSPDSLRDYFSKFGEIRECMVMRDPTTKRSRGFGFVTFADPASVDKVLGQP

HHELDSKTIDPKVAFPRRAQPKMVTRTKKIFVGGLSANTVVEDVKQYFEQFGKVEDAMLM

FDKTTNRHRGFGFVTFENEDVVEKVCEIHFHEINNKMVECKKAQPKEVMFPPGTRGRARG

LPYTMDAFMLGMGMLGYPNFVATYGRGYPGFAPSYGYQFPGFPAAAYGPVAAAAVAAARG

SVLNSYSAQPNFGAPASPAGSNPARPGGFPGANSPGPVADLYGPASQDSGVGNYISAASP

QPGSGFGHGIAGPLIATAFTNGYH

>Homo_sapiens.GRCh38.pep.all.fa_ENSP00000284073.2

MEANGSQGTSGSANDSQHDPGKMFIGGLSWQTSPDSLRDYFSKFGEIRECMVMRDPTTKR

SRGFGFVTFADPASVDKVLGQPHHELDSKTIDPKVAFPRRAQPKMVTRTKKIFVGGLSAN

TVVEDVKQYFEQFGKVEDAMLMFDKTTNRHRGFGFVTFENEDVVEKVCEIHFHEINNKMV

ECKKAQPKEVMFPPGTRGRARGLPYTMDAFMLGMGMLGYPNFVATYGRGYPGFAPSYGYQ

FPGFPAAAYGPVAAAAVAAARGSGSNPARPGGFPGANSPGPVADLYGPASQDSGVGNYIS

AASPQPGSGFGHGIAGPLIATAFTNGYH

>Homo_sapiens.GRCh38.pep.all.fa_ENSP00000313616.3

MADLTSVLTSVMFSPSSKMFIGGLSWQTSPDSLRDYFSKFGEIRECMVMRDPTTKRSRGF

GFVTFADPASVDKVLGQPHHELDSKTIDPKVAFPRRAQPKMVTRTKKIFVGGLSANTVVE

DVKQYFEQFGKVEDAMLMFDKTTNRHRGFGFVTFENEDVVEKVCEIHFHEINNKMVECKK

AQPKEVMFPPGTRGRARGLPYTMDAFMLGMGMLGYPNFVATYGRGYPGFAPSYGYQFPDY

LPVSQDIIFIN

>Homo_sapiens.GRCh38.pep.all.fa_ENSP00000462914.1

MADLTSVLTSVMFSPSSKMFIGGLSWQTSPDSLRDYFSKFGEIRECMVMRDPTTKRSRGF

GFVTFADPASVDKVLGQPHHELDSKTIDPKVAFPRRAQPKMVTRTKKIFVGGLSANTVVE

DVKQYFEQFGKVSAGSPGLS

>Homo_sapiens.GRCh38.pep.all.fa_ENSP00000392607.2

MMRGYIKKVLKNTSYHVNQYLFIQHGWSGSGDQSVLYSCNLMRMVTRTKKIFVGGLSANT

VVEDVKQYFEQFGKVEDAMLMFDKTTNRHRGFGFVTFENEDVVEKVCEIHFHEINNKMVE

CKKAQPKEVMFPPGTRGRARGLPYTMDAFMLGMGMLGYPNFVATYGRGYPGFAPSYGYQF

PGFPAAAYGPVAAAAVAAARGSGSNPARPGGFPGANSPGPVADLYGPASQDSGVGNYISA

ASPQPGSGFGHGIAGPLIATAFTNGYH

>Homo_sapiens.GRCh38.pep.all.fa_ENSP00000462264.1

MVTRTKKIFVGGLSANTVVEDVKQYFEQFGKVEDAMLMFDKTTNRHRGFGFVTFENEDVV

EKVCEIHFHEINNKMVECKKAQPKEVMFPPGTRGRARGLPYTMDAFMLGMGMLGYPNFVA

TYGRGYPGFAPSYGYQFPDYLPVSQDIIFIN

>Mus_musculus.GRCm38.pep.all.fa_ENSMUSP00000103542.1

MEANGSPGTSGSANDSQHDPGKMFIGGLSWQTSPDSLRDYFSKFGEIRECMVMRDPTTKR

SRGFGFVTFADPASVDKVLGQPHHELDSKTIDPKVAFPRRAQPKMVTRTKKIFVGGLSAN

TVVEDVKQYFEQFGKVEDAMLMFDKTTNRHRGFGFVTFENEDVVEKVCEIHFHEINNKMV

ECKKAQPKEVMFPPGTRGRARGLPYTMDAFMLGMGMLGYPNFVATYGRGYPGFAPSYGYQ

FPGFPAAAYGPVAAAAVAAARGSGSNPARPGGFPGANSPGPVADLYGPASQDSGVGNYIS

AASPQPGSGFGHGIAGPLIATAFTNGYH

>Mus_musculus.GRCm38.pep.all.fa_ENSMUSP00000090470.5

MEANGSPGTSGSANDSQHDPGKMFIGGLSWQTSPDSLRDYFSKFGEIRECMVMRDPTTKR

SRGFGFVTFADPASVDKVLGQPHHELDSKTIDPKVAFPRRAQPKMVTRTKKIFVGGLSAN

TVVEDVKQYFEQFGKVEDAMLMFDKTTNRHRGFGFVTFENEDVVEKVCEIHFHEINNKMV

ECKKAQPKEVMFPPGTRGRARGLPYTMDAFMLGMGMLGYPNFVATYGRGYPGFAPSYGYQ

FPGFPAAAYGPVAAAAVAAARGSVLNSYSAQPNFGAPASPAGSNPARPGGFPGANSPGPV

ADLYGPASQDSGVGNYISAASPQPGSGFGHGIAGPLIATAFTNGYH

>Mus_musculus.GRCm38.pep.all.fa_ENSMUSP00000119684.1

MFIGGLSWQTSPDSLRDYFSKFGEIRECMVMRDPTTKRSRGFGFVTFADPASVDKVLGQP

HHELDSKTIDPKVAFPRRAQPKMVTRTKKIFVGGLSANTVVEDVKQYFEQFGK

>Mus_musculus.GRCm38.pep.all.fa_ENSMUSP00000117497.1

MGKLFVGGLDWSTTQETLRSYFSQYGEVVDCVIMKDKTTNQSRGFGFVKFKDPNCVGTVL

ASRPHTLDGRNIDPKPCTPRGMQPERTRPKEGWKGPRSDSSKSNKIFVGGIPHNCGETEL

REYFKKFGVVTEVVMIYDAEKQRPRGFGFITFEDEQSVDQAVNMHFHDIMG

>Mus_musculus.GRCm38.pep.all.fa_ENSMUSP00000101001.1

MNSAGADEIGKLFVGGLDWSTTQETLRSYFSQYGEVVDCVIMKDKTTNQSRGFGFVKFKD

PNCVGTVLASRPHTLDGRNIDPKPCTPRGMQPERTRPKEGWKGPRSDSSKSNKIFVGGIP

HNCGETELREYFKKFGVVTEVVMIYDAEKQRPRGFGFITFEDEQSVDQAVNMHFHDIMGK

KVEVKRAEPRDSKNQAPGQPGASQWGSRVAPSAANGWAGQPPPTWQQGYGPQGMWVPAGQ

AIGGYGPPPAGRGAPPPPPPFTSYIVSTPPGGFPPPQGFPQGYGAPPQFSFGYGPPPPPP

DQFAPPGVPPPPATPGAAPLAFPPPPSQAAPDMSKPPTAQPDFPYGQYGYGQDLSGFGQG

FSDPSQQPPSYGGPSVPGSGGPPAGGSGFGRGQNHNVQGFHPYRR

>Mus_musculus.GRCm38.pep.all.fa_ENSMUSP00000101000.3

MNSAGADEIGKLFVGGLDWSTTQETLRSYFSQYGEVVDCVIMKDKTTNQSRGFGFVKFKD

PNCVGTVLASRPHTLDGRNIDPKPCTPRGMQPERTRPKEGWKGPRSDSSKSNKIFVGGIP

HNCGETELREYFKKFGVVTEVVMIYDAEKQRPRGFGFITFEDEQSVDQAVNMHFHDIMGK

KVEVKRAEPRDSKNQAPGQPGASQWGSRVAPSAANGWAGQPPPTWQQGYGPQGMWVPAGQ

AIGGYGPPPAGRGAPPPPPPFTSYIVSTPPGGFPPPQGFPQGYGAPPQFSFGYGPPPPPP

DQFAPPGVPPPPATPGAAPLAFPPPPSQAAPDMSKPPTAQPDFPYGQYAGYGQDLSGFGQ

GFSDPSQQPPSYGGPSVPGSGGPPAGGSGFGRGQNHNVQGFHPYRR

>Mus_musculus.GRCm38.pep.all.fa_ENSMUSP00000089958.5

MNSAGADEIGKLFVGGLDWSTTQETLRSYFSQYGEVVDCVIMKDKTTNQSRGFGFVKFKD

PNCVGTVLASRPHTLDGRNIDPKPCTPRGMQPERTRPKEGWQKGPRSDSSKSNKIFVGGI

PHNCGETELREYFKKFGVVTEVVMIYDAEKQRPRGFGFITFEDEQSVDQAVNMHFHDIMG

KKVEVKRAEPRDSKNQAPGQPGASQWGSRVAPSAANGWAGQPPPTWQQGYGPQGMWVPAG

QAIGGYGPPPAGRGAPPPPPPFTSYIVSTPPGGFPPPQGFPQGYGAPPQFSFGYGPPPPP

PDQFAPPGVPPPPATPGAAPLAFPPPPSQAAPDMSKPPTAQPDFPYGQYGYGQDLSGFGQ

GFSDPSQQPPSYGGPSVPGSGGPPAGGSGFGRGQNHNVQGFHPYRR

>Mus_musculus.GRCm38.pep.all.fa_ENSMUSP00000120516.1

METDAPQPGLASPDSPHDPCKMFIGGLSWQTTQEGLREYFGQFGEVKECLVMRDPLTKRS

RGFGFVTFMDQAGVDKVLAQSRHELDSKTIDPKVAFPRRAQPKMVTRTKKIFVGGLSVNT

TVEDVKHYFEQFGKVDDAMLMFDKTTNRHRGFGFVTFESEDIVEKVCEIHFHEINNKMVE

CKKAQPKEVMSPTGSARGRSRVMPYGMDAFMLGIGMLGYPGFQATTYASRSYTGLAPGYT

YQFPEFRVERSPLPSAPVLPELTAIPLTAYGPMAAAAAAAAVVRGTGSHPWTMAPPPGST

PSRTGGFLGTTSPGPMAELYGAANQDSGVSSYISAASPAPSTGFGHSLGGPLIATAFTNG

YH

>Mus_musculus.GRCm38.pep.all.fa_ENSMUSP00000070415.5

XSWQTTQEGLREYFGQFGEVKECLVMRDPLTKRSRGFGFVTFMDQAGVDKVLAQSRHELD

SKTIDPKVAFPRRAQPKMVTRTKKIFVGGLSVNTTVEDVKHYFEQFGKVDDAMLMFDKTT

NRHRGFGFVTFESEDIVEKVCEIHFHEINNKMVECKKAQPKEVMSPTGSARGRSRVMPYG

MDAFMLGIGMLGYPGFQATTYASRSYTGLAPGYTYQFPEFRVERSPLPSAPVLPELTAIP

LTAYGPMAAAAAAAAVVRGTGSTPSRTGGFLGTTSPGPMAELYGAANQDSGVSSYISAAS

PAPSTGFGHSLGGPLIATAFTNGYH

>Mus_musculus.GRCm38.pep.all.fa_ENSMUSP00000144032.1

XECVVTEGLREYFGQFGEVKECLVMRDPLTKRSRGFGFVTFMDQAGVDKVLAQSRHELDS

KTIDPKVAFPRRAQPKMVTRTKKIFVGGLSVNTTVEDVKHYFEQFGKVDDAMLMFDKTTN

RHRGFGFVTFESEDIVEKVCEIHFHEINNKMVECKKAQPKEVMSPTGSARGRSRVMPYGM

DAFMLGIGMLGYPGFQATTYASRSYTGLAPGYTYQFPAIPLTAYGPMAAAAAAAAVVRGT

GSTPSRTGGFLGTTSPGPMAELYGAANQD

>Mus_musculus.GRCm38.pep.all.fa_ENSMUSP00000143900.1

MVTRTKKIFVGGLSVNTTVEDVKHYFEQFGKVDDAMLMFDKTTNRHRGFGFVTFESEDIV

EKVCEIHFHEINNKMVECKKAQPKEVMSPTGSARGRSRVMPYGMDAFMLGIGMLGYPGFQ

ATTYASRSYTGLAPGYTYQFPAIPLTAYGPMAAAAAAAAVVRGTGSHPWTMAPPPGSTPS

RTGGFLGTTSPGPMAE

>Osativa_323_v7.0.protein_primaryTranscriptOnly.fa_LOC_Os10g33230.1

MESDQGKLFIGGISWETTEEKLRDHFAAYGDVSQAAVMRDKLTGRPRGFGFVVFSDPSSVDAALVDPHTLDGRTVDVKRA

LSREEQQAAKAANPSAGGRHASGGGGGGGGAGGGGGGGGGDAGGARTKKIFVGGLPSNLTEDEFRQYFQTYGVVTDVVVM

YDQNTQRPRGFGFITFDAEDAVDRVLHKTFHDLSGKMVEVKRALPREANPGSGSGGRSMGGGGGGYQSNNGPNSNSGGYD

SRGDASRYGQAQQGSGGYPGYGAGGYGAGTVGYGYGHANPGTAYGNYGAGGFGGVPAGYGGHYGNPNAPGSGYQGGPPGA

NRGPWGGQAPSGYGTGSYGGNAGYAAWNNSSAGGNAPTSQAAGAGTGYGSQGYGYGGYGGDASYGNHGGYGGYGGRGDGA

GNPAAGGGSGYGAGYGSGNGGSGYPNAWADPSQGGGFGASVNGVSEGQSNYGSGYGGVQPRVAQ*

>Osativa_323_v7.0.protein_primaryTranscriptOnly.fa_LOC_Os07g39560.3

MEADSGKLFVGGISWETDEDRLREYFSRFGEVTEAVIMRDRNTGRARGFGFVVFTDAGVAERVTMDKHMIDGRMVEAKKA

VPRDDQSITSKNNGSSIGSPGPGRTRKIFVGGLASNVTEVEFRRYFEQFGVITDVVVMYDHNTQRPRGFGFITYDSEDAV

DKALHKNFHELNGKMVEVKRAVPKEQSPGPAARSPAGGQNYAMSRVHSFLNGFNQGYNPNPIGGYGMRVDGRYGLLTGAR

NGFSSFGPGYGMGMNSESGMNANFGANSSFVNNSNGRQIGSFYNGSSNRLGSPIGYVGLNDDSGSLLSSMSRNVWGNENL

NYPNNPTNMSSFAPSGTGGQMGITSDGINWGGPTPGHGMGNISSLGLANLGRGAGDSFGLPSGSYGRSNATGTIGEPFSA

PPNAYEVNNADTYGSSSIYGDSTWRFTSSEIDMPPFGNDLGNVDPDIKSNIPASYMGNYTVNNNQTSRGITS*

>Osativa_323_v7.0.protein_primaryTranscriptOnly.fa_LOC_Os02g12850.1

MAGYPEENQHGMNGYEEEEEVEEVEGFDEEGRPGRRGGGGRDGGDGGAYGDASGDDGRAPGGDSSGKIFVGGVAWETTEE

SFSKHFEKYGAITDSVIMKDKHTKMPRGFGFVTFSDPSVIDKVLEDEHVIDGRTVEVKRTVPREEMSSKDGPKTRKIFVG

GLPSSLTEDELREHFSPYGKIVEHQIMLDHSTGRSRGFGFVTFESEDSVERVISEGRMRDLGGKQVEIKKAEPKKHGGDH

SSNGRSSHGSGGGYRSSYRSGGAAASGGGGGGGGGGSGSSGGYGYGAGYRSAGGGYYDSTAYGYGRGGYGYGGNAGFGSG

YGGGYGGSLYGGAYGAYGAYGGGAYGGGAYGGGAYGGGAYGGAPGAYGGAGGYGSYGGAGAGGAGGRGSSRYHPYGK*

>Osativa_323_v7.0.protein_primaryTranscriptOnly.fa_LOC_Os06g37000.2

MKDKHTKMPRGFGFVTFSDPSVIDKVLQDEHTIDGRTVEVKRTVPREEMSSKDGPKTRKIFVGGIPPSLTEDKLKEHFSS

YGKVVEHQIMLDHGTGRSRGFGFVTFENEDAVERVMSEGRMHDLAGKQVEIKKAEPKKPGGGDSSSNGRHSHGSGGGHRS

SYRGSGGGNSGSSSSGGYGGYGGGYRSAAAAYYGSTGYAGYGRGYGYGGNPAFGSGFGSGYGGSMYGGPYGAYGAYGGAY

GGGGAYGAPGGYGAGGYGAYGGAGGMGGGGSTSGRGSSRYHPYGK*

>Osativa_323_v7.0.protein_primaryTranscriptOnly.fa_LOC_Os11g41890.1

MEADAGKLFIGGISWDTNEDRLREYFDKYGEVVEAVIMRDRATGRARGFGFIVFADPAVAERVIMEKHMIDGRMVEAKKA

VPRDDQHALSKSGGSAHGSPGPSRTKKIFVGGLASTVTEADFRKYFEQFGTITDVVVMYDHNTQRPRGFGFITYDSEDAV

DKALFKTFHELNGKMVEVKRAVPKELSPGPSMRSPVGGFNYAVNRANNFLNGYTQGYNPSPVGGYGMRMDARFGLLSGGR

SSYPSFGGGYGVGMNFDPGMNPAIGGSSSFNNSLQYGRQLNPYYSGNSGRYNSNVSYGGVNDSTGSVFNSLARNLWGNSG

LSYSSNSASSNSFMSSANGGLGGIGNNNVNWGNPPVPAQGANAGPGYGSGNFGYGSSETNFGLGTNAYGRNAGSGVVNTF

NQSTNGYGRNFGDSSGGGGGGGGGSIYGDTTWRSGSSELDGTSPFGYGLGNAASDVTAKNSAGYMGH*

>Ppatens_318_v3.3.protein_primaryTranscriptOnly.fa_Pp3c26_4470V3.1.p

MEGGGDVERMQEVAPDLNGTLEDNDVDDEEQRQLDAVEDDEPHPDDEEERNNNGFQNGGSARDSMVESGFKDSPRDSQGK

ASGSGGKIFIGGLSWDTSTDNLQSHFKKYGEIIDAVIMKDRSTGHPRGFGFVTFADPAVCDNVVLDKHVIDGRTVEAKKS

VPRENMAASKGPKTKKIFVGGIPPSITDEEFKSYFASFGSVVEHQIMQDHSTGRSRGFGFVTFDSEQVVEDILAHGKMHE

LGGKQVEIKKAEPKRPIQDSPGGGYPGGGYPGGRGGGYGAGGVGGYAGYDGYGGSGSAGYGASYRSGGYGGRGGGYGGGY

GGGYGGGYGYGGGYGGGLGVGSYGSGVGGGYGGSLGYGSGGGYGSGMMGGYGDGADGYGSMGYGGYSGTGGYGAGYSSGG

GYGGSYGSSRGYSSSSGGGGSSRGRYHPYGRS*

>Ppatens_318_v3.3.protein_primaryTranscriptOnly.fa_Pp3c2_9350V3.1.p

MDSDQGKLFIGGISWETTEEKLRDYFKSYGEVVETVIMKDRATGRARGFGFVIFADPSVADRVAGEKHTIDGRTVEAKKA

VPRDEQQNMPRSNSVGGLGPATSHTRTKKIFVGGLASTVTEDDFKKYFEQFGTITDVVVMYDHNTQRPRGFGFITFDSED

AVENVIQKSFHELNEKMVEVKKAIPKELSGGSGRAAAGFGGGGRGGGNYGSGYSQGGYGGGSPGGGSGYGGGSPGGGSGY

GGRGDSRYGPSPGGRGGYPGYGAAGYGGAGFGSSGGYGAGMNGGYGGSAYGSSGGYGSGAGGYGSAGYGNAPGAGGYGSA

AGYGSGGVGSAGYGSAPAGGRSAWGSGGGGYGTGSGSTGYGGSGTGSMSGYGNAGWGTTGTGAGAAVSSGYGSAGYGYGS

SDAGYGSTGGYGGRGSGYGSSAGGYGTAGAGAGSGSYGGGYGDAYSGYADNTWRSSGTDPYSSGGTGSVGGGTGGGFGVG

GGASADGAVNGASGYGGGYGVAGRQTQRGPDARFRPYPAAGDRTG*

>Ppatens_318_v3.3.protein_primaryTranscriptOnly.fa_Pp3c2_9200V3.1.p

MDSDQGKLFIGGISWETTEEKLRDYFKSYGEVVETVIMKDRATGRARGFGFVIFADPSVADRVAGEKHTIDGRTVEAKKA

VPRDEQQNMPRSNSVGGQGPATSHTRTKKIFVGGLASTVTEDDFKKYFEQFGTITDVVVMYDHNTQRPRGFGFITFDSED

AVENVIQKSFHELNEKMVEVKKAIPKELSGGSGRAAAGFGGGGRGGGIYGSGYSQGGYGGGSPGGESGYGGGSPGGGSGY

GGRGGYPGYGAAGYGGAGFGSSGGHGAGMNGGYGGSAYGSSGGYGSGAGGYGSAGYGNAPGAGGYGSAAGYGSGGVGSAG

AWGSGGGGYGTGSGSTGYGGSGTGSMSGYGNAGWGTTGTGAGAAVSSGYGSAGYGYGSSDAGYGSTGGYGGRGSGYGSSA

GGYGTAGAGAGSGSYGGGYGDAYSGYADNTWRSSGTDSYSSGGTGSVGGGTGGGFGVGGGASADGAVNVASGYGGGYGVA

GRQTQRGVST*

>Ppatens_318_v3.3.protein_primaryTranscriptOnly.fa_Pp3c1_33600V3.1.p

MDSDQGKLFIGGISWETTEEKLRDYFKSYGEVAETVIMKDRATGRARGFGFVIFADPSVADRVAGEKHTIDGRTVEAKKA

VPRDEQQSIPRSNSVGGQGPATSHTRTKKIFVGGLASTVTEDDFKKYFEQFGTITDVVVMYDHNTQRPRGFGFITFDSED

AVENVIQKSFHELNDKMVEVKKAIPKELSAGSGRAGGGFGGGGRGGGNYGSGYSQGGYGGGSPGGGSGYGGRGDSRYGPT

PVGRGGYPGYGAAGYGAAGFGSSAGYGAGMNGGYGGSAYGSGGGYGSGAGGYGSAGYGSAPGAGGYGSGAGYGSGGTGSA

GYGSAPAGGRSAWGSGGGGYGTGASSTGYGGSGSGSMSGYGNAAWGTTGTGAGAAASSGYGSAAGYGYGSSETGYGSAGG

YGGRGSGYGSSGGGGGYGSSGAGTGSGSYGGGGYGDAYSGYADNTWRSSGSEAYNSGTTGSVGGGSGAGFGVGGGASAEG

AVNX

>Ppatens_318_v3.3.protein_primaryTranscriptOnly.fa_Pp3c1_33540V3.1.p

MDSDQGKLFIGGISWETTEEKLRDYFKSYGEVAETVIMKDRATGRARGFGFVIFADPSVADRVAGEKHTIDGRTVEAKKA

VPRDEQQSIPRSNSVGGQGPATSHTRTKKIFVGGLASTVTEDDFKKYFEQFGTITDVVVMYDHNTQRPRGFGFITFDSED

AVENVIQKSFHELNDKMVEVKKAIPKELSAGSGRAGGGFGGGGRGGGNYGSGYSQGGYGGGSPGGGSGYGGRGDSRYGPT

PVGRGXX

>Pvulgaris_442_v2.1.protein_primaryTranscriptOnly.fa_Phvul.009G074200.1.p

MNLKMDSDRAKLFVGGISRETTEDVLKFHFAKYGAVLDSTISLDHSTRCPRGFGFVTFSDLSAADQALQDTHVILGRTVE

VKKAIPRSEQQQHQNQLYSRGGGNYSNSCDCGSDTNVRTKKIFVGGLPAGISEEEFKQYFERFGRITDVVVMQDSVTHRP

RGFGFITFDSEESVQSVMVRSFHDLNGRQVEVKRAVPKEGNHGYDGFSKLRSKSERGVPKSFAPYSPRNMLPGFAPLPWY

SSDGIYSYGSNAQGCWYPMGGYGGNGYAIPSDASRNFWYGPMMACPQACQVPYASAVPNLAYFGGRIGIVGSGVGSWGYG

GILGSGTNFKFDQPFIGNGFVPGNTTLSWCKAECCGLFEFERKHW*

>Pvulgaris_442_v2.1.protein_primaryTranscriptOnly.fa_Phvul.008G286500.2.p

MPFLIPYDKMQSDNGKLFIGGISWDTNEERLREYFSTYGEVVEAVIMKDRTTGRARGFGFVVFSDPAVAEIVIKEKHNID

GRMVEAKKAVPRDDQNILNRNSGSIHGSPGPGRTRKIFVGGLASTVTESDFKKYFDQFGTITDVVVMYDHNTQRPRGFGF

ITYDSEEAVDKVLLKTFHELNGKMVEVKRAVPKELSPGPSRTPLGGYNYGLSRVNSFLNGFTQGYSPSSLGGYGLRADGR

FSPVASGRSGFAPFGSGYGMSMNFEPGLNAGFGGNANFNSNLSYGRGVNPYFIGSSNRFGSPVGFESGNGGNNSFFSSVT

RNLWGNGGLSYGTNSANSNAYIGSGSGSIGGNTFGNTGVNWGGSSAISGQGGGNNMSQSSGNLGYGSGDNNYGLGTGGYG

RNTGTAFAPTSSYSASNGGVDGAFADFYNNSSVYGDPTWRSSNSERDGSGPFGYGLGGAASDVSTKNSPGYVGGYTVNKR

QPNRGIAT*

>Pvulgaris_442_v2.1.protein_primaryTranscriptOnly.fa_Phvul.008G158124.1.p

MDSDQGKLFIGGISWDTTEDKLKEHFGNYGDVLSTSVMREKNTGKPRGFGFVVFADPTILDRVLEDKHVIDGRTVDAKKA

FSREDQQISVTSRGGNSNSGMNSGNGGNIRTKKIFVGGLPPTLTEEKFRQYFESYGHVTDVVVMYDQNTGRPRGFGFISF

DTEEAVDRVLHKSFHDLNGKQVEVKRALPKDANPGASGRMMGGAGGGGSGIGGYQGYGASGGNQNAYDGRMDSSRYMQPQ

SAAGGFPPYGSSAYSAPGYGYGPANNGIGYGAYGSYGGATAGYGGPAGATYGNPNVPNAAYAGGPPGGPRSSWPAQAPSG

YGSMGYGNTAPWGAPSGGAGSGGGGPGSATAGQSPGGAAGYGSQGYGYGGYGGYGGSDSSYNSSAYGTVGGRTGSAPNNS

ASGPGGGELTSSGGSGSYMSGGYGDANGNSGYGNAAWRSEQTHASGNYGTPQGNGGQVGYGGGYGGAQSRQAQQQ*

>Pvulgaris_442_v2.1.protein_primaryTranscriptOnly.fa_Phvul.008G274900.1.p

MANWAWLVAEQMESDLGKLFIGGISWDTDEERLREYFGKYGEVIEAVIMRDRTTGRARGFGFVVFSDPVVAERVIMDKHI

IDGRTVEAKKAVPRDDQITVNRQQTGGIQGSPGPGRTKKIFVGGLPSTITESDFKKYFDQFGTITDVVVMYDHNTQRPRG

FGFITYDSEEAVDRVLYKTFHELNGKMVEVKRAVPKELSPGPSRSPLVGYNYGLNRTSSFLNSYAQGFNMSPIGNYGVKM

EGRFSPLTTARSGFTPFGSNGYGMGVNLDSGLNPSYGGTSGYGGSLGYGRISPLYNGNPNRYTTPIGNNGGSGRTDSLMN

SASRSVWGNGGLNSAANNPASPGTYLGSGSGAFGVSIGNSGANWGSSVPTQGGGAASGYSTWGNTYEGGDNSIGLGGGGY

GRNSSPSVPQSSTFTAPTGGYEGSYGDLYRSGSVYSDSTWQSAASEMDASGSFGYGGLGGMASDDPVKSSEGFIGNYNVI

SRQTNRGIAA*

>Pvulgaris_442_v2.1.protein_primaryTranscriptOnly.fa_Phvul.006G053000.1.p

MESDLGKLFIGGISWDTDEERLKEYFGKYGEVIEAVIMRDRVTGRARGFGFVVFADPSVAERVIMDKHIIDGRTVEAKKA

VPRDDQLNINRQQQSGSAHASPGPGRTKKIFVGGLPSTITESDFKKYFDQFGTITDVVVMYDHNTQRPRGFGFITYDSEE

AVDRVLYKTFHELNGKMVEVKRAVPKELSPGPNRSPLIGYNYGLNRASSYLNSYAQGYSMSPIGGYGVRMDGRFSPLTGG

RSGFTPFGNTGYGMGMNLDSGLSPTFGGTSNYGSNLGYGRLLNPFYNGNSGRYTTPIGYSGVNGRSDSLMNSPSRNLWGN

GGLNNASNPISPSPFLGSGSGAFGVSIGNSGTGWGPSIPAQGGGAASGYGTGNNIYEGGDSSFGLGGAGYGRNNSSGVTP

ASTFNVSTGGYEGSYGDLYRTGSGSVYNDAAWRSAVSEIDGSGSFGYGLGGIASDDPVKNSEGYIGNYNVTSRQPNRGIA

A*

>Saccharomyces_cerevisiae.R64-1-1.pep.all.fa_YOL123W_mRNA

MSSDEEDFNDIYGDDKPTTTEEVKKEEEQNKAGSGTSQLDQLAALQALSSSLNKLNNPNS

NNSSSNNSNQDTSSSKQDGTANDKEGSNEDTKNEKKQESATSANANANASSAGPSGLPWE

QLQQTMSQFQQPSSQSPPQQQVTQTKEERSKADLSKESCKMFIGGLNWDTTEDNLREYFG

KYGTVTDLKIMKDPATGRSRGFGFLSFEKPSSVDEVVKTQHILDGKVIDPKRAIPRDEQD

KTGKIFVGGIGPDVRPKEFEEFFSQWGTIIDAQLMLDKDTGQSRGFGFVTYDSADAVDRV

CQNKFIDFKDRKIEIKRAEPRHMQQKSSNNGGNNGGNNMNRRGGNFGNQGDFNQMYQNPM

MGGYNPMMNPQAMTDYYQKMQEYYQQMQKQTGMDYTQMYQQQMQQMAMMMPGFAMPPNAM

TLNQPQQDSNATQGSPAPSDSDNNKSNDVQTIGNTSNTDSGSPPLNLPNGPKGPSQYNDD

HNSGYGYNRDRGDRDRNDRDRDYNHRSGGNHRRNGRGGRGGYNRRNNGYHPYNR

>Sbicolor_454_v3.1.1.protein_primaryTranscriptOnly.fa_Sobic.010G171100.1.p

MAGYGEENQNAMNGYEEEEEEEEVEEVEEVYEEEEEEGDGDGEGEGEADEGAAAAEAAAMAAETRSVVGDRGLAEAAGNV

DAGGEDGRDADSGGGDSSGKIFVGGVAWETTEETFTKHFQKYGAITDSVIMKDKHTRMPRGFGFVTFSDPSVLDRVLEDA

HVIDGRTVEVKRTVPKEEMSSKDGPKTKKIFVGGIPSSLTEDKLKEHFSSYGKVVEHQIMLDHSTGRSRGFGFVTFESED

AVERVMSEGRMHDLGGKQVEIKKAEPKKPGGGDSSSNGRHSRGGGHRDSYRGSGGGSGVSGNSSGGGYGYGGGYRSTAAA

YYGSTAYGAYGRGYGYGGTAAGYGSGYGSAYGGSMYGGPYGAYGAYGGAYGGGAYGAPGGYGAGGYDSYGGAGGMGGGGS

AGGRGSSRYHPYGK*

>Sbicolor_454_v3.1.1.protein_primaryTranscriptOnly.fa_Sobic.005G197000.1.p

MEADAMKLFIGGISWDTNEDHLLKYFKKYGEVEEAIVMRDRATGRSRGFGFIMFADPAVAKHVIMEKHMIDGRMVEAKKA

IARDDHHSLNNIHGSAHGLQRPKHRKKIFVGGLASNVTKEDVIKHFKQFGTIIDVVVVYDHHTQRPRGFGFITYDSEDAV

HRALIKTFQKLKGKMVEVKRAIRKEPSPIPSMCSPINGFNYVTGRANSFLNGYTQHYNMSPLGGYGMRMDECFGLLSGGN

TNNGYPSFGGSHGIGMNFDTSMNPCTESGSSFNSGIQHGWHLNPYYNGNLGGFNNSVSYGGVSGNNGLLFDSLVHNLWDI

FGLNHSSSSTSSNFFVPFGNGDLSGIGNNNVNWGNPHHVPAPGANNCSVYCTGNFCYGSSETNFGQGSSVYGRNIGSSGD

NNLNQSTSGYARNFGDSSVGGGSIYGDTTWTSRSFEHDVTKMNSTGHMGH*

>Sbicolor_454_v3.1.1.protein_primaryTranscriptOnly.fa_Sobic.004G095300.1.p

MAGYPEDNQHALNGYDEEEVDEEEGHPGRRGGRDGASGYGDAGGEDGRGAGGDSSGKIFVGGVAWETTEESFSKHFEKYG

AITDSVIMKDKHTKMPRGFGFVTFSDPSVIDKVLEDDHVIDGRTVEVKRTVPREEMTTKDGPKTRKIFIGGLPPSLTEDE

LKDHFSSYGKVVEHQIMLDHSTGRSRGFGFVTFESEDSVERVISEGRMRDLGGKQVEIKKAEPKKHGSDHSSNGRSSHGG

GGYRSSYRSGGGAGNASSGSSGGGGGGYGYGGAYRSAAAGYGYDGGAGAGYGYGRGYGYGGNAGFGSGFGGGYGGSMYGG

AYGAYGAYGGGAYGGGAYGGGAYGGGAYGGAPGGYGTGGYGGYGGAGGASGGTGGGSTGARGSSRYHPYGK*

>Sbicolor_454_v3.1.1.protein_primaryTranscriptOnly.fa_Sobic.001G222900.1.p

MESDQGKLFIGGISWETTEEKLSEHFSAYGEVTQAAVMRDKITGRPRGFGFVVFADPAVVDRALQDPHTLDGRTVDVKRA

LSREEQQASKAANPSGGRNTGGAGGGGGGGAGDASGARTKKIFVGGLPSTLTEDGFRQYFQTFGSVTDVVVMYDQNTQRP

RGFGFITFDSEDAVDRVLHKTFHDLGGKLVEVKRALPREANPGGSSGGRSGGSGGYQSNNGHNTSSGSYDGRSDGGRYGQ

AQQGSGGYPGYGAGGYGAGAAGYGYGANPAVGYGNYGAGGYGGVPAAYGGHYGNPGAAGSGYQGGPPGSNRGPWGSQAPS

AYGTGGYGSSAGYSAWNNSSGGGNAPSSQAPGGPAGYGSQGYGYGGYGGDPSYASHGGYGAYGGRSDGAGNPATGGASGY

SAGYGSGGANSGYSSAWSDPSQGGGFGGSVNGGAEGQSNYGTGYGSVQPRVAQ*

>Sbicolor_454_v3.1.1.protein_primaryTranscriptOnly.fa_Sobic.002G067100.1.p

MEAAAAAAVADYGKLFVGGISWETSEDRLRDYFGRFGEVTEAVIMRDRSTGRARGFGFVVFADPAVAERVTMDKHMIDGR

MVEAKKAVPRDDHSIVSKSNASSIGSPGPGRTRKIFVGGLPSNVTEADFRRYFEQFGVITDVVVMYDHNTQRPRGFGFIT

YDSEDAVDKALHKSFHELNGKMVEVKRAVPKEQSPGPVVRSPVGVGQNYAMNRVHSFLNGFNQGYNPNPIGGYGMRVDGR

YGLLSGARNGFSSFGPGYGMGMNVEGGMSGTFGASSGFISNSNGRQMGSYFNASSNRLGSPMGYLGLNDVPGSMLSSMSR

NVWGNGSLNYPSNPTNMSAFSSPGNGGQVGITGDSWGGLPSAHGMGSISSLGSGNLGRGAGDNTFGLPSGSYGRSNSTGT

IGEPFSASGNTYEVNNPDTYGSSSIYGGTAWRFASSEVDMPSFGHDIGNVDPTIK*

>Sbicolor_454_v3.1.1.protein_primaryTranscriptOnly.fa_Sobic.002G067600.1.p

MEAAAAVADYGKLFVGGISWETSEDRLRDYFGRFGEVTEAVIMRDRSTGRARGFGFVVFADPAVAERVTMDKHMIDGRMV

EAKKAVPRDDHSIVSKSNASSIGSPGPGRTRKIFVGGLPSNVTEADFRRYFEQFGVITDVVVMYDHNTQRPRGFGFITYD

SEDAVDKALHKSFHELNGKMVEVKRAVPKEQSPGPVVRSPVGVGQNYAMNRVHSFLNGFNQGYNPNPIGGYGMRVDGRYG

LLSGARNGFSSFGPGYGMGMNVEGGMSGTFGASSGFISNSNGRQMGSYFNASSNRLGSPMGYLGLNDVPGSMLSSMSRNV

WGNGSLNYPSNPTNMNAFASPGNGGQVGITGDSWGGLPSAHGMGSISSLGSGNLGRGAGDNTFGLPSGSYGRSNSTGTIG

EPFSASVNTYEVNNPDTYGSSSIYGGTAWRFASSEVDMPSFGHDIGNVDPTIK*

>Sbicolor_454_v3.1.1.protein_primaryTranscriptOnly.fa_Sobic.002G355300.1.p

MEADSGKLFVGGISWETDEDRLREYFGRFGEVTEAVIMRDRNTGRARGFGFVVFADSGIAERVTMDKHMIDGRMVEAKKA

VPRDDQSIASKNNGSSIGSPGPGRTRKIFVGGLASNVTEVEFRRYFEQFGVITDVVVMYDHNTQRPRGFGFITYDSEDAV

DKALHKNFHELNGKMVEVKRAVPKEQSPGPIARSPVGGQNYAMSRVHNFLNGFNQGYNPNPLGGYGMRVDGRYGLLTGAR

NGFSSFGPGYGMGMNVEGGMSGYFGASSGFVNSSNGRQIGSYFNGSSNRLGSPIGYVGLNDDSGSILSSMSRNVWGNGNL

NYTGNPTNTNSFAPPGSGGGIPGDEISWGSLTSAHGMGNISSLGSGNLGRGTGDHNFGLPSGNYVRSNSTGTIGEPFSAS

TNAYESNNPGAYGSSSIYGDSTWRFNSSEVDMPPFGHDLGNVDPDIKSEISASYMGNYTVNNNQTSRGITS*
